# Older age is associated with a shift from ventral to dorsal PCC in pathways connecting the DMN with visual and limbic areas and a directionality shift in pathways connecting the DMN with sensorimotor areas

**DOI:** 10.1101/2022.12.04.519066

**Authors:** Gadi Goelman, Rotem Dan, Ondrej Bezdicek, Robert Jech, Dana Ekstein

**Author notes:** **<u>Corresponding Author:</u>** Gadi Goelman, PhD, Department of Neurology, Hadassah Hebrew University Medical Center, Jerusalem 91120, Israel.

## Abstract

Alterations in the default mode network (DMN) are known to associated with aging and with neurological and psychiatric diseases. We assessed age-dependent changes in interactions within the DMN and between the DMN with other brain areas, and correlations of these interactions with a battery of neuropsychological tests to formulate a macroscopic model of ageing.

Using a novel multivariate analysis method, enabling determination of directed pathways, on resting-state functional MRI data from fifty young and thirty-one old healthy individuals, we identified directed intra- and inter-DMN directed pathways that differed between the groups and used their correlations with neuropsychological tests to infer their behavioral meaning.

Pathways connecting the DMN with visual and limbic regions in old subjects engaged at BOLD low frequency and involved the dorsal PCC. In young subjects, they were at high frequency and involved the ventral PCC. Pathways combining the sensorimotor (SM) and the DMN, were efferent in young subjects (DMN<SM) and afferent in old subjects (DMN>SM). Most inter-DMN pathways in the old subjects, correlated with reduced speed and working memory and, were DMN efferent (DMN>).

We suggest a macroscopic model of aging centered in the DMN. It suggests that the reduced sensorimotor efferent, probably brought about from reduced physical activity, and the increased need to control such activities by the mPFC, causes a higher dependency on external versus internal cues. This results in a shift from ventral to dorsal PCC of inter-DMN pathways. Consequently, the model combines physical activity with ageing.

## 1. Introduction

Cognitive decline, reduced processing speed, difficulties with attention and with working and episodic memory are common findings associated with healthy aging [1-5]. Studies using multiple connectivity measures of healthy young and old subjects have shown reduced default mode network (DMN) connectivity and an altered functional organization of many primary sensory and cognitive networks in elderly subjects [1]. The DMN is best known for being active during wakeful rest, such as daydreaming or mind-wandering. It is also active when an individual is thinking about others, thinking about themselves, remembering the past, and planning the future [6]. Changes in the DMN have been shown to be associated with neurological and psychiatric diseases, e.g. it is vulnerable to the early stages of Alzheimer disease (AD) pathology [7, 8].

Within the DMN, the posterior cingulate cortex (PCC) has been shown to be functionally segregated, with its ventral part primarily connected to other DMN regions and supporting internally directed thought [9]. Its dorsal part is primarily connected to multiple networks including the sensorimotor network and supports externally directed attention [10]. Subjects with AD and with amnestic mild cognitive impairment (aMCI) have been shown to have a modulated PCC connectivity. While the ventral PCC network was atrophied in aMCI and AD patients, the dorsal network was only atrophied in AD patients [11]. It is, therefore, of great clinical relevance to understand the functionality of the dorsal and ventral PCC in elderly subjects.

Connectivity of the DMN has been addressed in many resting-state functional magnetic resonance imaging (rs-fMRI) studies in young and old subjects however, with inconsistent findings. Some have shown reduced anterior-posterior intra DMN connectivity [12], while others have revealed a complex increase and decrease in the connectivity of specific connections [13-17]. We suggest that this inconsistency may be attributed to the methods used to assess connectivity, and especially to the use of bivariate and undirected connectivity. Consider, for example, the connectivity between the PCC and the supplementary motor area. If these regions are involved in two separated information flows such as: mPFC→PCC→SMA and SMA→PCC→Angular, the bivariate undirected connectivity between the PCC and the SMA gives an average coupling which is an inaccurate estimation of the functional coupling between these regions.

To derive a more precise connectivity estimation, we recently introduced a multivariate analysis application that can be used with rs-fMRI data. This method identifies directed interactions among four different anatomical regions, i.e. it enables one to define functional, directed couplings relating to anterograde information flow (i.e., pathways) among four different anatomical regions [18-22]. Computer simulations of the Kuramoto model have tested the accuracy of this analysis [18], and its applications to the human brain were demonstrated in the above sited studies.

Hereby, we hypothesize that DMN connectivity is age-dependent and reflect the reduced cognitive and adaptive behavior abilities of old subjects. To test these, we identified functional directed pathways within the DMN and between the DMN and other brain areas that differed between 50 healthy young and 31 healthy old participants, and calculated their associations with a battery of neuropsychological tests to infer their behavioral meaning.

## 2. Materials and Methods

### 2.1 Subjects

This study used data from young and old healthy participants who underwent fMRI measurements at two different sites using similar systems and identical protocols. The study was approved by the Hadassah Medical Center, Jerusalem, Israel, Ethics Committee and the Ethical Committee of the General University Hospital in Prague, Czech Republic. All participants provided written informed consent prior to inclusion in the study, which was carried out in compliance with the Declaration of Helsinki.

#### The young subject group

Fifty-two, healthy, young, undergraduate students at the Hebrew University of Jerusalem, Israel were recruited for this study. To exclude past or present psychiatric disorders, participants were evaluated by a clinical psychologist or a psychiatrist using the Structured Clinical Interview for DSM-IV (SCID-5-CV). Additional exclusion criteria were neurological disorders, and, for women, the use of hormonal contraceptives, pregnancy or breastfeeding. Two male subjects were excluded by these criteria, yielding a final sample of 20 men (age:23.9±2.9 years) and 30 women (23.9±2.4 years). Part of these data was used in a previous study [23].

#### The old subject group

40 elderly subjects were recruited from a community in Prague. Nine of them were excluded due to: severe atrophy or vascular lesions (n=5), in-scanner motion (n=3), or use of lithium (n=1) which yielded a final sample of 31 subjects. The final group included 16 females (age 61.2±6.4 years) and 15 males (65.2±8.8 years). Exclusion criteria were a history of psychotic symptoms, depression, dementia or a cognitive state on the Montreal Cognitive Assessment < 22. Psychomotor speed and working memory, executive function, language, long-term memory and visuospatial function were measured (see below). Part of these data was used in previous studies [21, 24].

### 2.2 Neuropsychological tests of old subjects

Old subjects underwent neuropsychological assessment by an experienced neuropsychologist during a preliminary visit approximately two weeks before the MRI session. A battery of tests that included two neuropsychological tests for each of the following five domains was used: (1) Psychomotor speed and working memory: Trail Making Test, part A [25] and Digit span backwards from the Wechsler Adult Intelligence Scale, third revision (WAIS-III) [26]; (2) Executive function: Tower of London [27] and semantic verbal fluency [28]; (3) Language: Boston Naming Test, Czech version [29, 30] and WAIS-III Similarities (Wechsler, 1997); (4) Long term memory: Rey Auditory Verbal Learning Test, delayed recall [31] and Brief Visuospatial Memory Test, revised, delayed recall [32, 33]; (5) Visuospatial function: CLOX [34] and Judgment of Line Orientation [35]. The score on each test was transformed into a z-score using the Rankit formula [36]. The z-scores of each domain were summarized for each subject to create summary scores, with a higher score indicating a better function.

Table 1 lists the participants’ demographic data and the neuropsychological evaluation tests.

**Table 1.** Demographic and memory characteristics of the subjects. RAVLT- Rey Auditory Verbal Learning Test, Delayed Recall, BVMT-R - Brief Visuospatial Memory Test, Revised, Delayed Recall.

| **Demographic and neuropsychological variables** | **Young**<br>**Mean ± SD (range)**<br>**(n=50)** | **Elderly**<br>**Mean ± SD (range)**<br>**(n=31)** |
| --- | --- | --- |
| Age, years | 23.9±2.6 (19-28) | 63.2±7.89 (46-83) |
| Gender (male/female) | 20/30 | 15/16 |
| Education, years | 14.1±1.75 (12-20) | 14.8±3.5 (11-25) |
| **Long-term memory** |  |  |
| RAVLT-30 | | 8.97±2.5 (2-12) |
| BVMT-R-30 | | 10.23±1.70 (5-12) |
| **Long-term term memory summary** | | 67.8±22.2 (8.0-94.2) |
| **Visuospatial function** |  |  |
| Royall's CLOX (CLOX I) | | 13.26±1.15 (11-15) |
| Judgment of Line Orientation | | 24.65±3.88 (11-29) |
| **Visuospatial function summary** | | 59.4±17.9 (23.8-86.7) |
| **Psychomotor speed and working memory** |  |  |
| Trail Making Test, part A | | 35.2±9.1 (20-58) |
| Digit Span Backwards | | 6.7±2.2 (4-12) |
| **Attention summary** | | 65.12±17.4 (35.9-92.3) |
| **Executive function** |  |  |
| Tower of London | | 26.1±4.1 (16-34) |
| Semantic fluency | | 65.6±11.8 (38-93) |
| **Executive function summary** | | 59.8±18.4 (14.4-93.1) |
| **Language** |  |  |
| Wechsler Adult Intelligence Scale-III Similarities | | 23.7±5.7(11-32) |
| Boston Naming Test (Czech version) | | 54.4±6.5 (27-60) |
| **Language summary** | | 62.0±13.1 (34.6-81.5)<br>57.4±25.6 (7-97.4) |

### 2.3 MRI data acquisition and preprocessing

MRI data of the young and old participants were acquired with 3T MR scanners (Magnetom Skyra, Siemens, Germany). The group of young participants was scanned at the neuroimaging center of the Hebrew University of Jerusalem, Israel; and the group of old participants was scanned at Charles University in Prague, Czech Republic. At both locations, participants underwent a 10-min resting-state fMRI (rs-fMRI) during fixation on a visual crosshair, using the same MRI acquisition protocols. Functional images were acquired using a T2*-weighted gradient-echo, echo-planar imaging sequence with TR=2 s, TE=30 ms, image matrix=64×64, field of view= 192 × 192 mm, flip angle=90°, resolution= 3 × 3 × 3 mm, interslice gap=0.45 mm. Each brain volume comprised 30 axial slices, and each functional run contained 300 image volumes. Anatomical images were acquired using a sagittal T1-weighted MP-RAGE sequence with TR=2.2 sec, TE=2.43 ms, resolution=1×1×1 mm. For the old participants group, T2-weighted images were collected as well, for diagnostic purposes, to exclude significant atrophy or any other pathological brain changes.

All functional MRI data underwent the following preprocessing using SPM12 (http://www.fil.ion.ucl.ac.uk/spm/software/spm12). Functional images were spatially realigned, coregistered to T1 anatomical images; slice-time corrected and normalized to MNI space. Further preprocessing was done in a CONN toolbox [37]. Potential confounding effects were regressed out using the aCompCor method for anatomical component-based noise correction [38]. These included: (i) outlier scans, i.e. censoring/scrubbing [39]. Outlier scans were identified based on the amount of subject motion in the scanner as measured by frame-wise displacement (FD) and global BOLD signals. Acquisitions with FD > 0.9mm or global BOLD signal changes > 5 standard deviations were considered outliers and removed by regression; (ii) the first 5 principal components (PCAs) of the CSF and white matter signals, to minimize the effects of physiological non-neuronal signals such as cardiac and respiratory signals; (iii) estimated subject-motion parameters and their first-order derivatives (a total of 12 parameters); (iv) session effects: the potential effects of the beginning of the session were removed by a step function convolved with the hemodynamic response function, in addition to the linear BOLD signal trend. After regression of all potential confounding effects, temporal band-pass filtering (0.008–0.09 Hz) was performed. Note that we did not apply global signal regression, due to its controversy. Global signal regression may introduce artifactual biases [40] and remove potentially meaningful neural components [41]. Instead, the aCompCor method [38] was applied to regress out the first 5 principal components (PCAs) of the CSF and white matter signals. This was done to minimize the effects of potential physiological non-neuronal signals such as cardiac and respiratory signals, without the risk of artificially introducing anticorrelations into the functional connectivity estimates.

## 3. Theory

### 3.1 Multivariate directed functional connectivity and correlation with behavior test

A detailed description of the analysis was presented in our previous publications [18-22], therefore, only the main points are summarized below. For a group of four weakly coupled BOLD temporal signals, with each signal corresponding to a different anatomical location, the analysis assumed that the phases contained temporal information of the order of their mutual coupling. This order is expressed in terms of specific relations between the four phases, as defined previously [18], and enables the definition of four-node pathways corresponding to information transfer among them. For resting-state data, we averaged over time, in the time-frequency wavelet space, to obtain frequency-dependent phase differences. We restricted the analysis to continuous, unidirectional pathways to define pathways among the four BOLD signals, i.e., pathways that started in one region and subsequently went through all the other regions. In this case, there were 24 possible pathways (listed in Supplementary Table 1). In this table, three of the four regions were predefined (symbolized by R1 to R3), while the fourth region (symbolized as “X”) was related to another brain region (see below). By choosing pathways that were invariant to the choice of reference-phase (see below), we guaranteed that all phase differences were below 2π [21, 22], thus obtaining unbiased pathways.

For each participant (*sub*), each pathway’s type (*k*) and each frequency (*ω*), a binary pathway value (*PW*) was defined as “1” for the cases where phase differences were in line with the pathway and “0” for when they were not (for detailed explanation and illustration see [21] and supplementary figure 1 their):

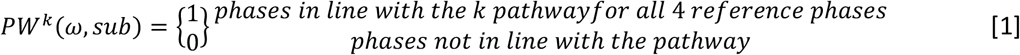

with ‘k’=1, 2… 24 corresponding to a pathway’s number in Supplementary Table 1, “ω” the frequency scale, and “*sub*” a subject.

A group pathway index (PWI) was defined as:

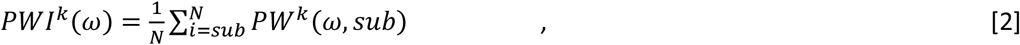

similar to the definition of the phase lag index (PLI) [42, 43] but describing the coherence among four regions, while PLI describes the coherence between two regions. We further note that averaging the wavelet coherences among participants solved the intrinsic time-frequency uncertainty [44, 45].

Comparisons between groups were performed as follows:

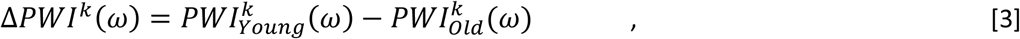

whose significance was obtained by non-parametric permutation tests.

To calculate the unique contribution of each neuropsychological domain, i.e., Long-term memory, Visuospatial function, Psychomotor speed and Working Memory, Executive function and Language (listed in Table 1) to the pathways, we used a partial correlation model between *PW*^*k*^(*ω, sub*) of Equation 1 and each neuropsychological domain, while statistically controlling for the contributions of all other neuropsychological domains as well as for age, gender, years of education and the frame-wise displacement (FD). Formally, we calculated the correlation between the residuals of *PW*^*k*^(*ω, sub*) of each neuropsychological domain. These residuals were obtained using linear regression with all other neuropsychological domains and with gender, age, education and FD. Multivariate wavelet calculations were performed with IDL version 8.2.0 (Exelis Visual Information Solutions, Inc.) using custom-developed software. The complex Morlet wavelet functions were chosen for wavelet analysis because they have been shown to provide a good trade-off between time and frequency localization [46]. We used 10 for the smallest scale, 2 for time resolution, and 21 scales to cover the entire frequency window. Wavelet software was provided by Torrence and Compo (http://paos.colorado.edu/research/wavelets) [44, 47] [44, 47] [44, 47] [44, 47] [44, 47] [44, 47] [44, 47]. For simplicity and easier evaluation, the 21 frequencies were further averaged into three frequency bands: low (average frequency 0.022 Hz, named scale 1), intermediate (average frequency 0.04 Hz, named scale 2), and high (average frequency 0.073 Hz, scale 3).

### 3.2 Experimental design

Owing to the large number of possible pathways where each fMRI voxel can be a region; we minimized the number of pathways as follows: (1) The Atlas of Intrinsic Connectivity of Homotopic Areas (AICHA)[48] with its 384 regions was used for region selection. The average BOLD signals from these regions for each subjects were used in the analysis. (2) Pathways were calculated with three preselected regions, while the fourth was any of the other 384 regions in the atlas. That is, instead of pathways with all the possible region’s combinations (384*C*_4_ = 8.9*E* + 8), we had 384 pathways for each of the three preselected regions. Note that this procedure enabled us to focus on a specific network by choosing the three preselected regions in that network.

### 3.3 Between-group pathways

Comparison between young and old participants was performed in four stages (see Figure 1 for illustration). The first stage aimed at identifying pathways that were significantly different between the groups. For that, we calculated four-node pathways with three predefined DMN regions. The three predefined regions were: one out of the two AICHA medial prefrontal cortex (mPFC) regions; one out of the three AICHA angular gyrus (Ang) regions; and one of the three AICHA posterior cingulate cortex (PCC) regions. The fourth region was any of the other AICHA regions (Figure 1A). Consequently, we calculated 18. (384 − 8) pathways with 18, the number of combinations of the predefined seeds. Specifically, we used the AICHA ‘G_Frontal_Med_Orb-1’ and ‘G_Frontal_Med_Orb_2’ as the two mPFC regions, the ‘G_Angullar_1’, ‘G_Angular_2’ and the ‘G_Angular_3’ as the three angular regions, and ‘G_Cingulum_Post_1’, ‘G_Cingulum_Post_2’ and ‘G_Cingulum_Post_3’ as the three PCC regions. Note that the ‘G_Cingulum_Post_1’ and the ‘G_Cingulum_Pos_2’ corresponded to the ventral PCC (vPCC) while the ‘G_Cingulum_Pos_3’ corresponded to the dorsal PCC (dPCC). All preselected regions were in the dominant (left) hemisphere. Pathways were calculated for each group using equation 2, and their differences were calculated by equation 3.

**Figure 1:**
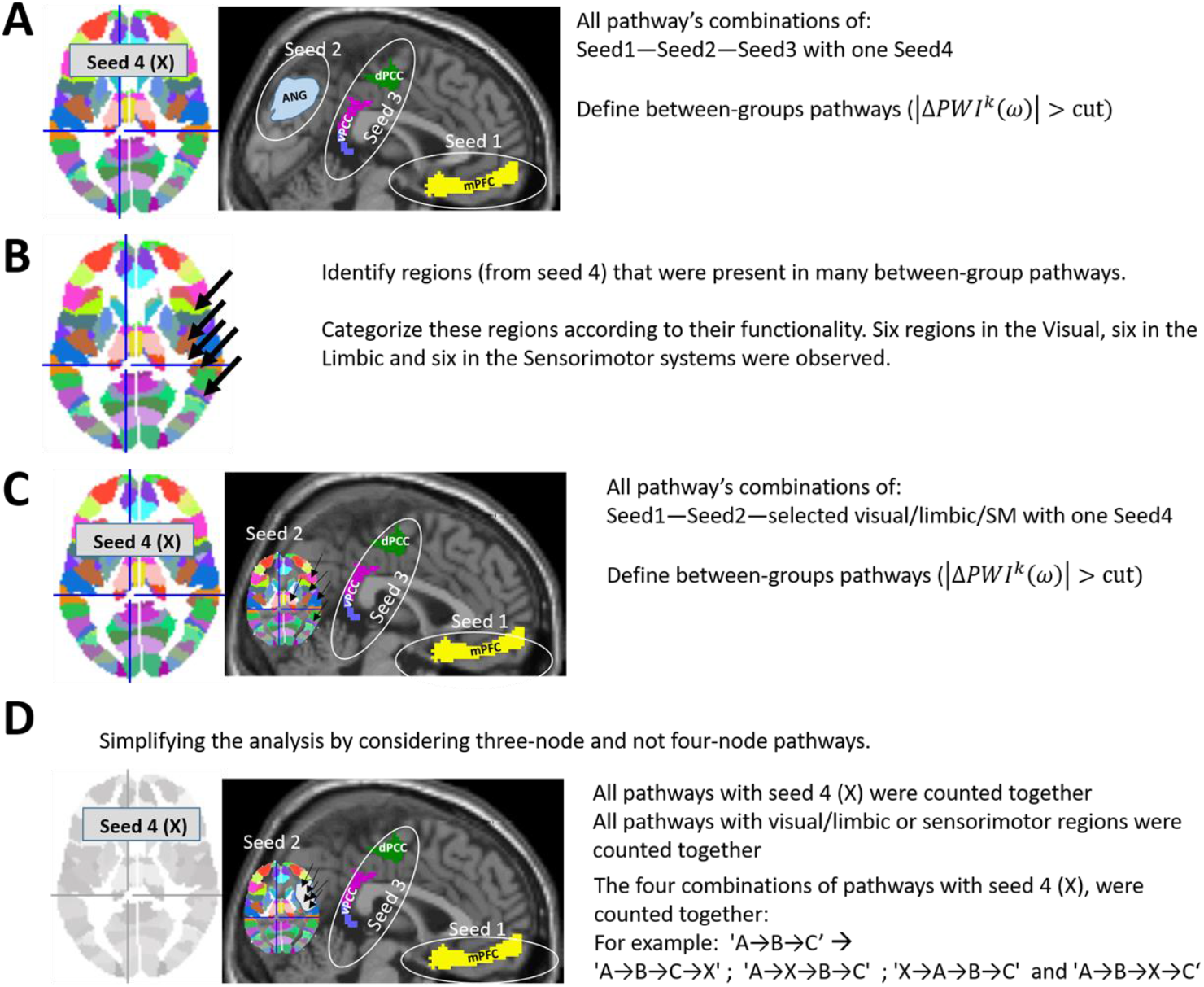
Illustration of the experimental design. **A**. Calculations of all between-group four-node pathway combinations with three predefined DMN regions and one brain region. All together 18. (384 − 8) between-group pathways were calculated. **B**. We identified 18 non-DMN regions that were in many between-group pathways: six from the visual, six from the limbic and six from the sensorimotor systems. **C**. Calculating all between-group four-node pathway combinations using two DMN predefined regions, one predefined region from the regions selected in B, and one free region. All together 36. (384 − 8) between-group pathways were calculated. **D**. To enable comparison between pathways, we calculated the occurrences of three-node between-group functional pathways (calculated in A and in C) by summing together the pathways of a different fourth seed and pathways with different fourth seed locations. These three-node pathways are presented in the text.

In the second stage, we aimed to identify extra-DMN regions involved in many pathways that were different between the two subject groups and to categorize them according to their systems (Figure 1B). Based on the highest occurrence at the end of the first stage that compared between the young and old groups, we selected 18 extra-DMN regions: six were in the visual system, six in the limbic system and six in the sensorimotor system.

In the third stage, we calculated the between-groups inter-DMN four-node pathways (Figure 1C). These were pathways with two predefined regions from the DMN, one from the selected regions of the visual, limbic or sensorimotor systems and one from all other brain regions. The preselected regions were a mPFC region and a PCC region from the DMN, and one from the six selected visual, limbic or sensorimotor regions. All together, we calculated 36. (84 − 11) pathways with 36 the number of combinations of the predefined regions.

In the fourth stage, we aimed to estimate which pathways differed the most between the groups. Since four-node pathways are binary, we calculated the occurrences (see below) of three-node, between groups, functional pathways (Figure 1D). In these calculations, pathways with two mPFC regions and pathways with two vPCC were summed together, as well as pathways within the six visual/limbic or sensorimotor regions.

### 3.4 Correlation with behavior

To identify the pathways in the old subject group that correlated with behavior, we used the above stages 3 and 4 to identify pathways that correlated with the five neuropsychological domains of Table 1. Specifically, we calculated the partial correlations of each neuropsychological domain with the inter-DMN four-node pathways of stage 3 while statistically controlling for the contributions of all other neuropsychological domains as well as for age, gender, FD and years of education. To enable comparison with the between-groups pathways and to simplify the analysis, we calculated the occurrences of three-node functional pathways, as in stage 4. We used binary assignments to define pathways, that is, pathways with a correlation value above a cutoff value were assigned ‘1’, pathways with a correlation value below minus the cutoff value were assigned ‘-1’, and all other pathways were assigned the value zero.

### 3.5 Pathway’s occurrence

To clarify the results, we reduced the number of pathways by deriving three-nodes from the four-node pathways, and by using the pathways’ occurrence. Specifically, we did not differentiate between pathways with different fourth region, and pathways with different fourth region’s location (recall that the fourth region was any non-preselected region). For example, the three-node pathway ‘A→B→C’, was defined by the sum of the following four-node pathways: ‘A→B→C→X’; ‘A→X→B→C’; ‘X→A→B→C’; and ‘A→B→X→C’ with X symbolized the forth regions that could be any of the AICHA regions. Consequently, the maximum number of a three-node pathway was:

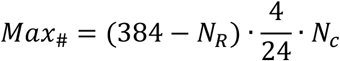

with 384 the number of AICHA regions, *N*_*R*_ the number of preselected regions, four the number of summed pathways, 24 the number of possible pathways and *N*_*c*_ the number of summed combinations. *N*_*R*_ equalled 8 for the intra-DMN, and equalled 11 for the inter-DMN pathways. *N*_*c*_ equalled 6 for the intra-DMN pathways (2 mPFC * 3 Ang) and 12 for the inter-DMN (2 mPFC * 6 regions). Note that averaging over the two vPCC regions was done in the last stage. Consequently, *Max*_#–*intra*_ = 376 and *Max*_#–*inter*_ = 746. The occurrence was presented in terms of percentiles of these maximum numbers.

### 3.6 Statistical analysis

We used permutation non-parametric tests to calculate the null distributions of Equations 2 and 3 for each frequency. Equation 2 was used within group while Equation 3 to make comparisons between groups. The null distributions were calculated using rs-fMRI signals from the mPFC, the angular gyrus, the PCC and the precuneus of the left hemisphere, using the automatic anatomical labelling (AAL) atlas [49]. We used a random number generator to select seeds from different participants to guarantee uncoupling. This process was repeated 10000 times and Equations 2 and 3 were calculated each time. As expected, the distributions of the 24 pathways were approximately identical. This is since, for null coupling, no differences between pathways were expected. The distributions implied that *PWI*^*k*^(*ω*_1_, *ω*_2_, *ω*_3_) and Δ*PWI*^*k*^(*ω*_1_, *ω*_2_, *ω*_3_) of Equations 2 and 3 for the young and the young-old groups when equaled to [0.125, 0.125, 0.33], corresponded to uncorrected p-value of ∼10^−3^. Similarly, these values for the old and the old-young groups when equaled, [0.17, 0.17 0.37], corresponded to uncorrected p-value of ∼10^−3^. Due to the high cutoff of pathways at the highest frequency, no results were obtained for this frequency scale (see below). For the partial correlation cutoff, we used the same uncorrected p-value (*p* = 10^−3^) which translated to |*R*| > 0.55. These uncorrected p-values matched to FDR (false discovery rate) corrected p-value of *α*_*FDR*_ = 0.05, corrected for multiple comparisons.

Supplementary Figure 1 illustrates the stages to obtain group and between-groups pathways.

### 3.7 Pathway’s directionality

To obtain directed pathways, we had to assume what the direction of coupling was, that is, whether the signal was transferred from right to left or from left to right, in the ranked pathways of Table 2. In other words, we had to determine whether a positive phase difference between regions *i* and *j* corresponded to the signal flow from *i* to *j* or from *j* to *i*. This factor, common to all coherent studies, depends on a reference phase which is an intrinsic factor of the system. Since our pathways were defined for a group, the reference phase must be equal for the all group’s data-sets which requires that all group’s data-sets acquired by the same system. Once this factor was assumed, we applied it to all pathways within a group because it is not specific to a pathway but common to all [43]. To infer directionality, we needed to identify a pathway whose directionality was known or could be assumed. Similar to our previous studies [18-22], we calculated thalamocortical pathways (using Equation 2), assuming that most pathways in the rs were bottom-up. For these calculations, the left thalamus, left primary motor and left primary sensory cortices were the preselected regions, focusing on the motor system of the dominant hemisphere, and the fourth region were rs-fMRI voxels using all 3D image voxels. Null distribution of these cases implied that the *PWI*^*k*^(*ω*_1_, *ω*_2_, *ω*_3_) of [0.2, 0.2, 0.39] corresponded to an uncorrected p value of *p*∼2 ∗ 10^−5^ and to corrected p∼0.001 with a cluster-size threshold of 100 [22]. We found that the majority of the (undirected) pathways were either “Thalamus-M1-S1-X” or “Thalamus-S1-M1-X” with “X” presenting the clusters in the motor system or frontal areas [21, 22]. Bottom-up processes suggest left-to-right directionality in the pathways of Table 2 in the old group and right to left directionality in the young group. This directionality was applied to all the pathways in this study.

**Table 2:**
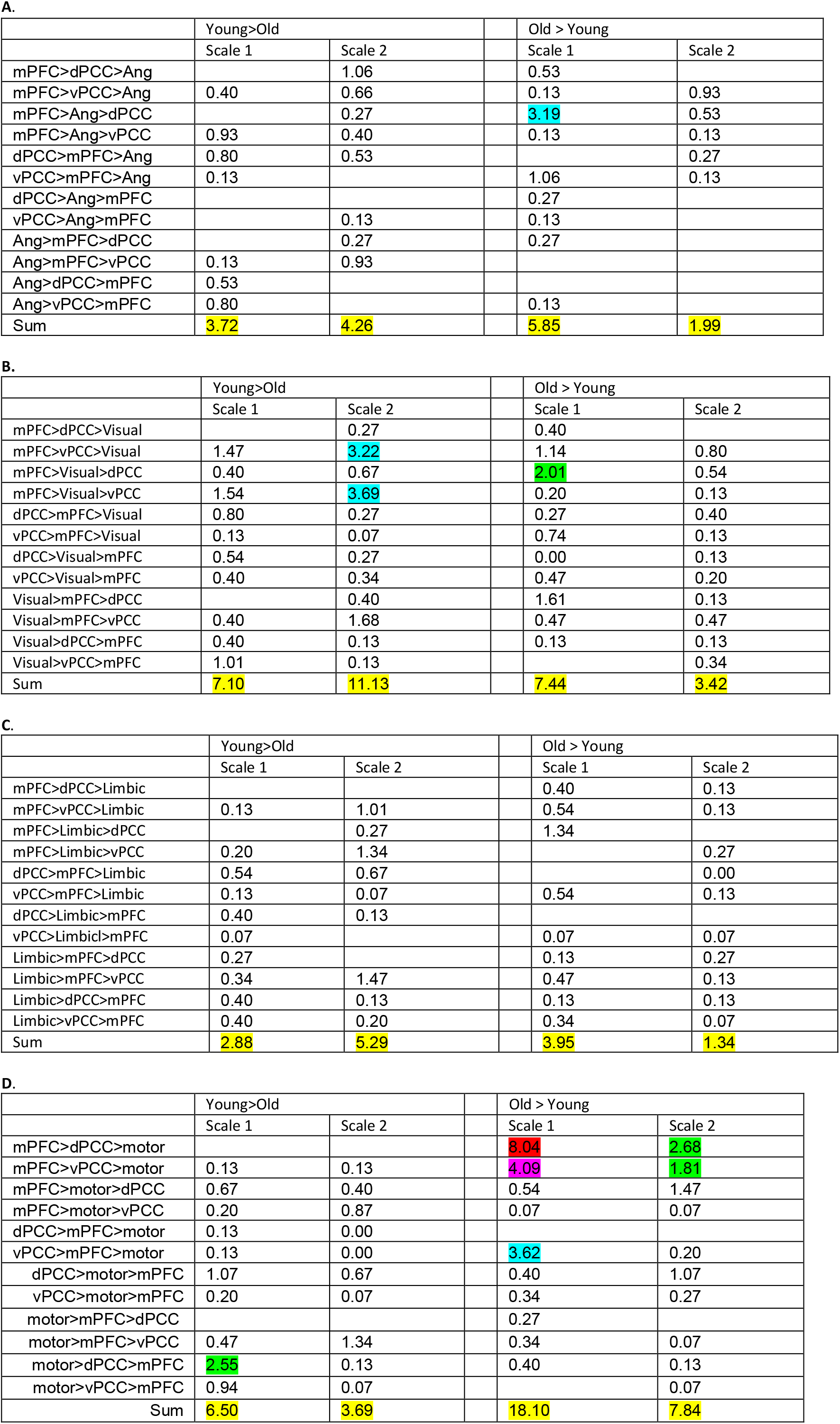
Percentages of 3-node between-group pathways (out of the maximum numbers). **A**. The 12 intra-DMN pathways. **B**. The 12 inter visual-DMN pathways. **C**. The 12 inter limbic-DMN pathways. **D**. The 12 inter sensorimotor-DMN pathways. Colors indicating decreasing occurrences red-purple-blue-and green.

### Data Availability

Data will be available upon request

## 4. Results

A three-node directed pathway has six permutations in the order of the nodes. Since we differentiated between the ventral and the dorsal PCC, we had 12 permutations. In the figures and tables below, we present the occurrences of the pathways represented by these 12 permutations for pathways within the DMN and pathways between it and regions outside the DMN. In addition, we present the pathways that correlated with the five neuropsychological domains tested in the old subject group. Note however, that the same three-node pathways could result from different four-node pathways. This is since the fourth node could be at different locations, and since each of the three nodes resembles one of several possible regions. For example, it can be one of the six visual regions or one of the three mPFC regions. For all the pathways, significant results were only obtained at the two lower scales of frequency. Below, we first describe the results of each group separately, then we show the pathways that differed between the age groups, and last we describe pathways that correlated with the five neuropsychological domains.

### 4.1 Inter-DMN pathways within each age group

Supplementary Table 2 depicts the one-group (young and old) occurrences of the 12 three-node directed pathways for frequency scales one and two for the pathways connecting the DMN with extra-DMN areas (refer to as ‘inter-DMN’ pathways, in percentage of the maximum number). Most pathways that connected the DMN with regions in the visual and the limbic systems of the young group included the ventral PCC (65% of the pathways connecting the DMN and the visual system and 45% of the pathways connecting the DMN and the limbic system). However, in the old subjects group, only 12% of the pathways connecting the DMN with the visual system, and 17% of the pathways connecting the DMN with the limbic system included the ventral PCC. No significant differences between the two age groups for pathways that included the dorsal PCC were observed. Furthermore, most of these pathways for the young group were at scale 2 (64% of the pathways connecting the DMN and visual areas and 68% of the pathways connecting the DMN and limbic areas). In contrast, most of the pathways of the old subject group were at scale 1 (69% of the pathways connecting the DMN with visual areas and 72% of the pathways connecting the DMN with limbic areas). The main difference between the groups for pathways connecting the DMN with sensorimotor regions was their directionality: 49% of the pathways were efferent to the mPFC in the old subject group, and 27% of the pathways were afferent to the mPFC in the young subject group.

### 4.2 Intra-DMN pathways between age groups

Table 2A gives the between-group occurrences of the 12 three-node directed pathways for frequency scales one and two for the intra-DMN pathways (in percentage of the maximum number). The numbers of pathways with positive (‘Young>Old’; implying stronger connectivity in young than in older subjects) and negative (‘Old>Young’; stronger connectivity in the older subjects) Δ*PWI*^*k*^(*ω*), were comparable. However, there were differences between the pathways in the two groups, both in their signal frequencies and in DMN nodes involved (Figure 2). While more Young>Old pathways were at higher frequency (scale 2; 56%), most of the Old>Young pathways were at the lowest frequency (scale 1; 75%). Furthermore, more Young>Old pathways were connected with the vPCC (56%), whereas most of the Old>Young pathways were connected with the dPCC (63%).

**Figure 2:**
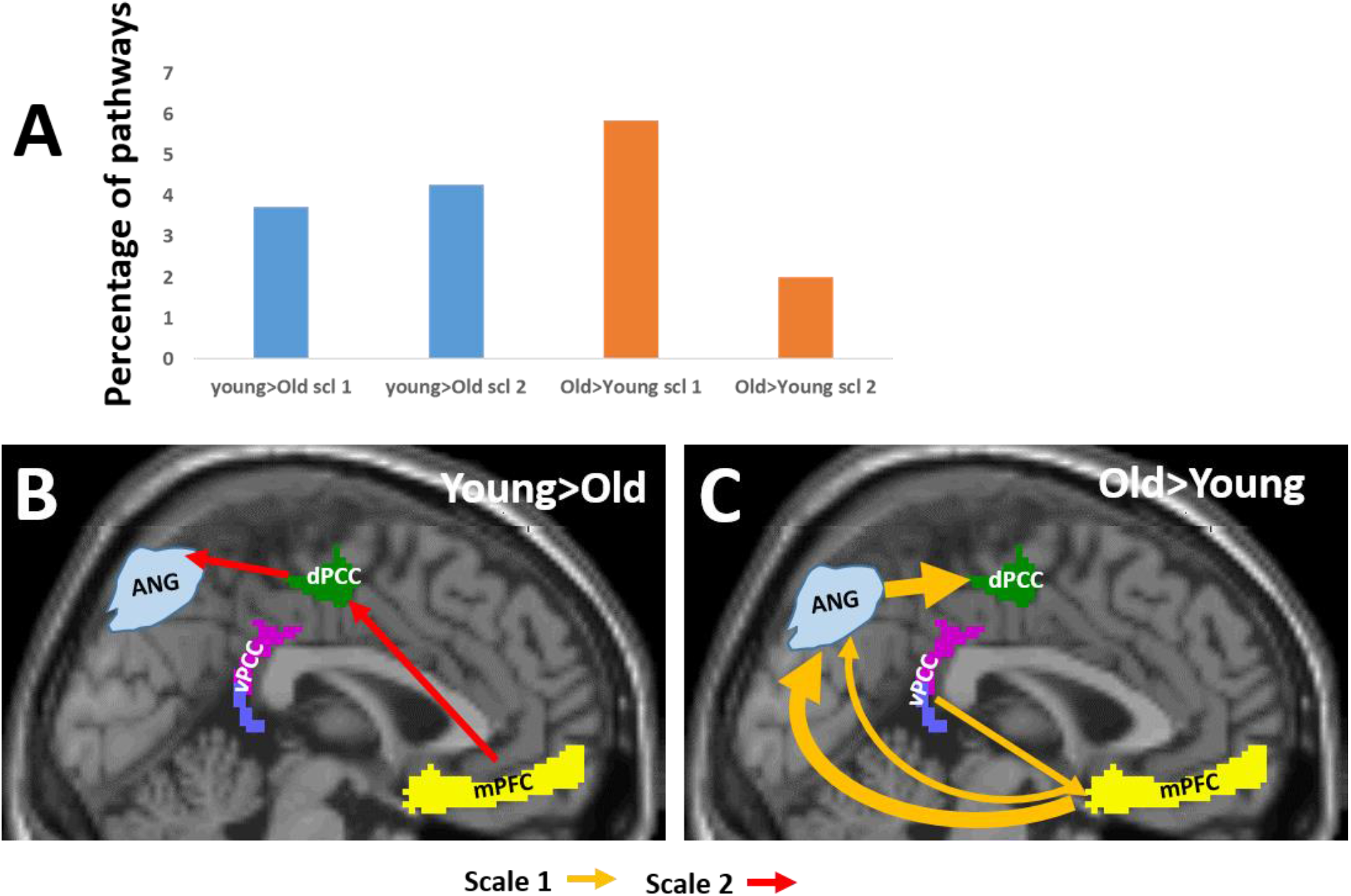
Three-node between-groups functional intra-DMN pathways. **A**. Percentiles of pathways (out of the maximum possible numbers) that were stronger in the young participants’ group (in blue) and in the old participants’ group (in orange) in scale 1 and scale 2. **B-C**. Illustrations of pathways that were stronger in the young participants (B) and were stronger in the old participants’ group (C), with pathways in light orange for scale 1 and in red for scale 2. ‘mPF’C is the medial prefrontal cortex, ‘vPCC’ and ‘dPCC’ are the ventral and dorsal posterior cingulate cortex and ‘ANG’ is the angular gyrus. Only pathways whose occurrences were >1% are shown with arrow width corresponding to pathway’s occurrences.

### 4.3 DMN-visual between group pathways

Table 2B gives the between-group occurrences of the 12 three-node directed pathways for frequency scales one and two for the pathways connecting the DMN with vision regions. These pathways involved mainly the ventral and dorsal precuneus and to a slighter extent the lingual, the lateral occipital gyri and the fusiform gyrus (Figure 3A). Specifically, the following six AICHA regions were found in the DMN-visual between group pathways: Precuneus 3, Precuneus 2, Precuneus 9, Fusiform, Occipital_lat 3 and Lingual 1. As shown in Table 2B, the number of pathways with positive (‘Young>Old’) Δ*PWI*^*k*^(*ω*) values was higher (63%) compared to the number of pathways with a negative values (‘Old>Young’). Furthermore, while 61% of the Young>Old pathways were at frequency scale 2, 68% of the Old>Young pathways were at scale 1 suggesting age dependent scale. In addition, 77% of the Young>Old pathways involved the vPCC, while the Old>Young pathways were about equal with both PCC sub-regions. Figure 3 illustrates these findings, showing the high occurrence pathways.

**Figure 3:**
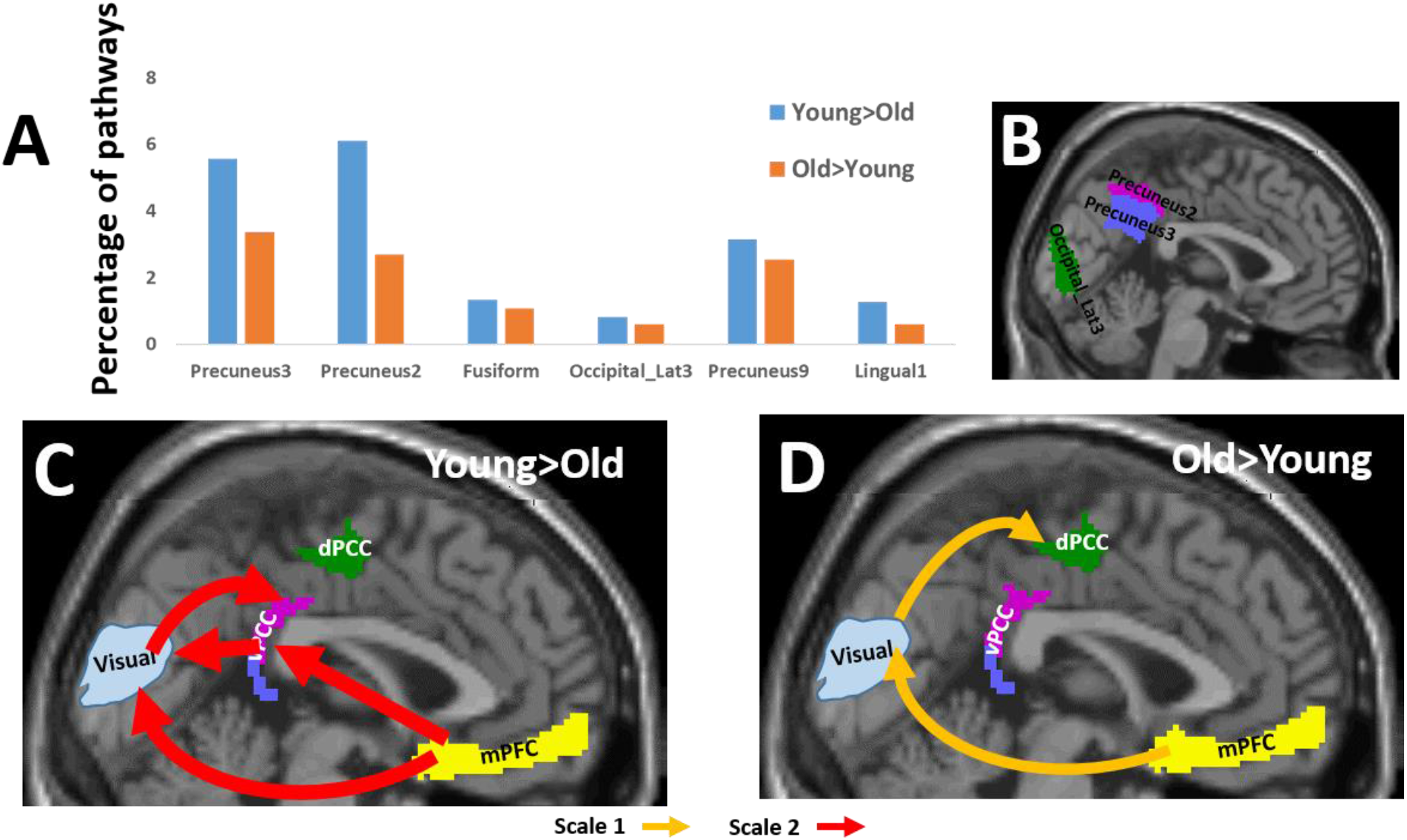
Three-node between-groups functional inter visual-DMN pathways. **A**. Percentiles of the pathways (scales 1 and 2 together) that include a visual region for the six regions. In blue, pathways that were stronger in the young participants’ group, and in orange, pathways that were stronger in the old participants’ group. **B**. A 2D sagittal projection of the three visual regions whose pathways differed most between the groups. **C-D**. Illustrations of the most dominant pathways that were stronger in the young participants (C) and were stronger in the old participants group (D), with pathways in light orange for scale 1, and in red for scale 2. Only pathways whose occurrences were >2% are shown with arrow width corresponding to pathway’s occurrences.

### 4.4 DMN-limbic between group pathways

Table 2C gives the between-group occurrences of the 12 three-node directed pathways for frequency scales one and two for the pathways connecting the DMN with limbic regions. DMN-limbic pathways that were different between the groups involved the medial parahippocampus, the adjacent hippocampus and, to a slighter extent, the putamen, the anterior insula and the caudate (Figure 4A). Specifically, the following six AICHA regions were found: Parahippocampus 1, Caudate 3, Parahippocampus 4, Anterior Insula 4, Putamen and Hippocampus 2. The number of pathways with positive Δ*PWI*^*k*^(*ω*) values was higher (61%) compared to the number of pathways with negative values (‘Old>Young’). Furthermore, while 65% of the Young>Old pathways were at scale 2, 75% of the Old>Young pathways were at frequency scale 1. In addition, 65% of the Young>Old pathways involved the vPCC, whereas the Old>Young pathways were about equal with both PCC sub-regions. Figure 4 illustrates these findings, showing the high occurrence pathways.

**Figure 4:**
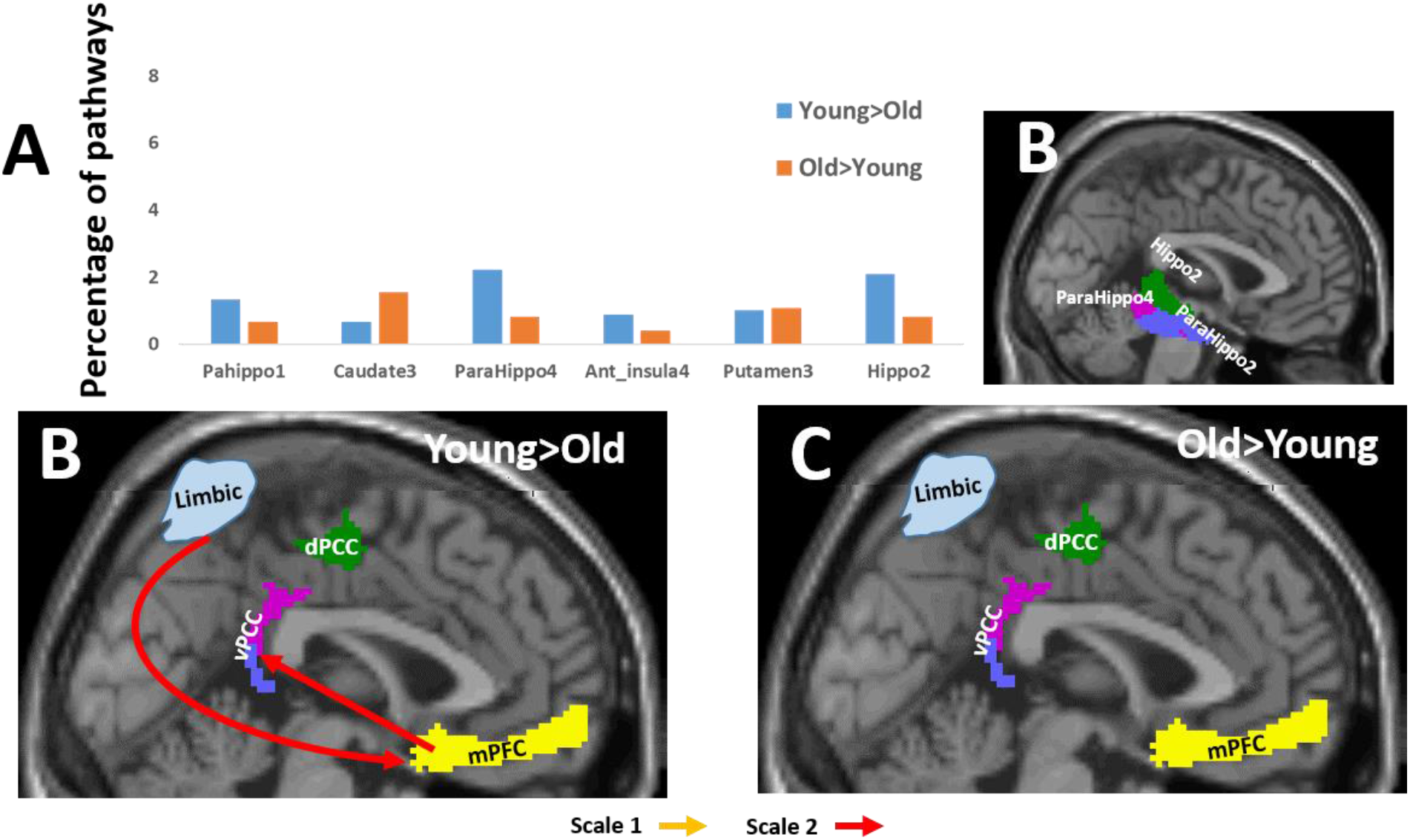
Three-node between groups functional inter limbic-DMN pathways. **A**. Percentiles of pathways (scales 1 and 2 together) that included a limbic region for the six regions. In blue, pathways that were stronger in the young participants’ group, and in orange, pathways that were stronger in the old participants’ group. **B**. A 2D sagittal projection of the three limbic regions whose pathways differed most between the groups. **C-D**. Illustrations of the most dominant pathways that were stronger in the young participants (C) and were stronger in the old participants group (D), with pathways in light orange for scale 1, and in red for scale 2. Since the occurrences of all pathways were low, only one whose occurrence is ∼1% is shown.

### 4.5 DMN-sensorimotor between group pathways

Table 2D gives the between-group occurrences of the 12 three-node directed pathways for frequency scales one and two for the pathways connecting the DMN with sesnsorimotor regions. Figure 5A shows that the inter DMN-sensorimotor pathways that differed between the groups involved mainly the postcentral, the ventral paracentral lobule, and the dorsal Rolando sulcus. Specifically, the following six AICHA regions were found: Paracentral lobule 4, Paracentral lobule 1, Rolando 3, Postcentral 3, Precentral 1 and Parietal Sub. Table 2D gives the between-group pathways’ occurrences for the inter DMN-sensorimotor pathways. The number of pathways with positive Δ*PWI*^*k*^(*ω*) values was lower (28%) compared to the number of pathways with negative value (‘Old>Young’). In both groups, most of the pathways were at low frequency scale (64% for Young>old and 70% for the Old>Young pathways) and they had comparable involvements of vPCC and dPCC (55% and 58% of dPCC in young>Old and Old>Young, respectively). However, the most significant difference between the groups was directionality: While 54% of the Young>Old pathways were efferent that is, lead from the sensorimotor system towards the DMN, only 5% were afferent that is in the opposite direction (while in the others the sensorimotor nodes were between the two DMN nodes). In contrast, 79% of the Old>Young pathways were afferent that is, started in the DMN and ended in the sensorimotor system with only 5% of the pathways efferent. Figure 5 illustrates these findings, showing the high occurrence pathways.

**Figure 5:**
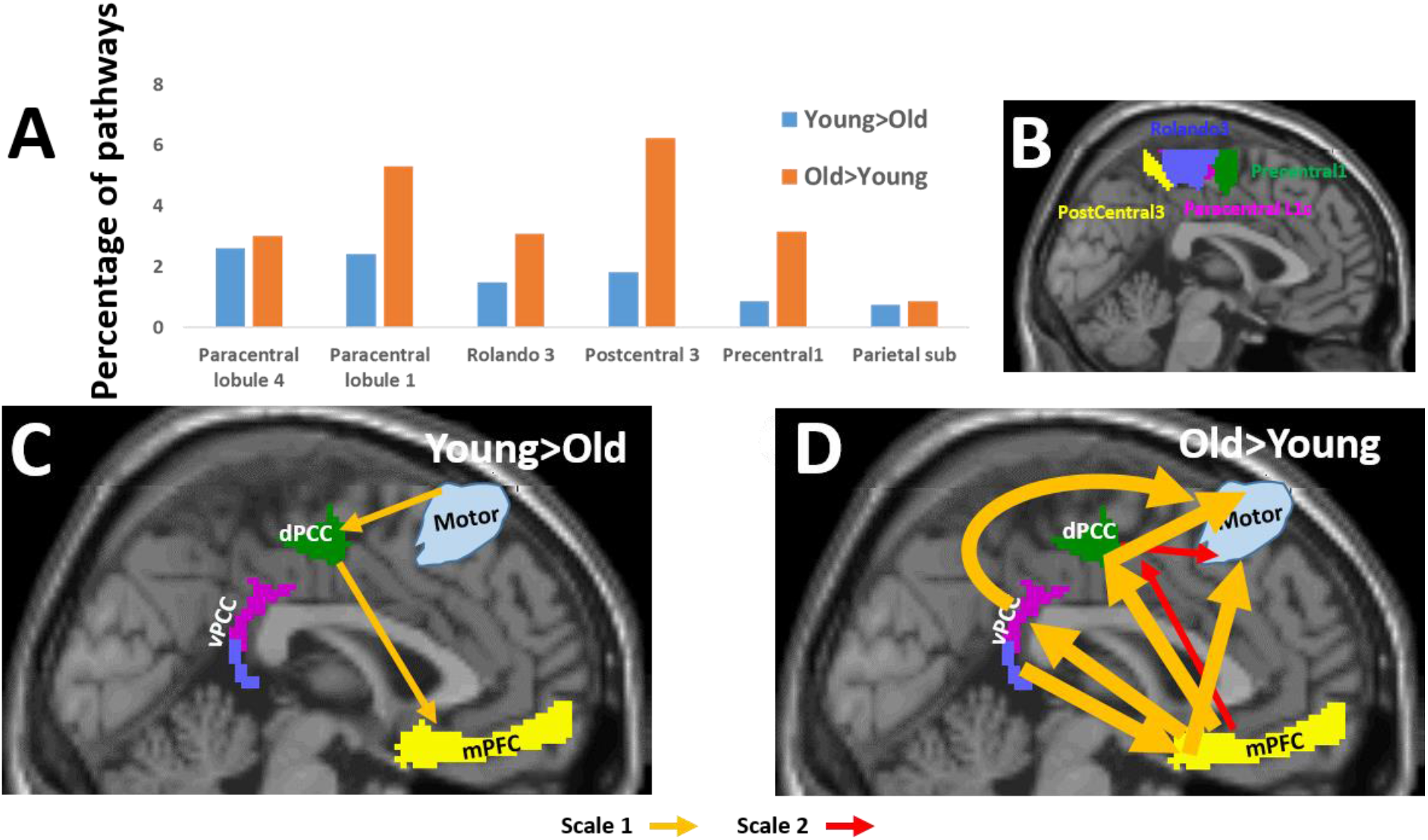
Three-node between groups functional inter sensorimotor-DMN pathways. **A**. Percentiles of pathways (scales 1 and 2 together) that included a sensorimotor region for the six regions. In blue, pathways that were stronger in the young participants’ group, and in orange, pathways that were stronger in the old participants’ group. **B**. A 2D sagittal projection of the three sensorimotor regions whose pathways differed most between the groups. **C-D**. Illustrations of the most dominant pathways that were stronger in the young participants (C) and were stronger in the old participants’ group (D), with pathways in light orange for scale 1, and in red for scale 2. Only pathways whose occurrences were >2% are shown with arrow width corresponding to pathway’s occurrences.

### 4.6 Pathways that correlated with neuropsychological domains

Supplementary Table 3 lists the occurrences of the 12 three-node directed pathways that were correlated with the five different neuropsychological domains, for the pathways connecting the DMN with extra-DMN brain regions (in percentage of the maximum number). Positive correlation implies an association between pathway’s indexes of Equation 1 and a better score, while negative correlation relates to an association between the pathway’s indexes and a lower score. For easier observation, we present in Figure 6 the numbers of pathways, for each frequency scale of these pathways, for the five neuropsychological domains. The figure shows that most of the pathways with negative correlation values were with the psychomotor speed and working memory (SWM) and with the visuospatial function (VSF) domains, while most of the pathways with positive correlation values were with the language and the memory domains. Figure 7 presents the occurrences of the pathways that negatively correlated with SWM and VSF (all scales summed together). It shows that reduced SWM was generally correlated with DMN efferent pathways (that is, leading from the DMN towards other extra-DMN brain areas, with p=0.039, one side t-test) in line with Supplementary Table 2 (t-test between the four ‘DMN>System’ pathways of all 3 scales versus the four ‘System>DMN’ of all three scales). Figure 8 presents the pathways that positively correlated with language and memory for all pathways connecting the DMN with other brain regions. It shows that most of the pathways that positively correlated with memory were efferent (that is, lead from the DMN towards one of the other regions with p=0.036, one side t-test). We also found a tendency for significance of the pathways that positively correlated with language to have an opposite directionality, i.e. from other brain regions toward the DMN (p=0.06, one side t-test). supplementary Figures 2-5 present the occurrences of pathways correlated with these four domains for all scales.

**Figure 6:**
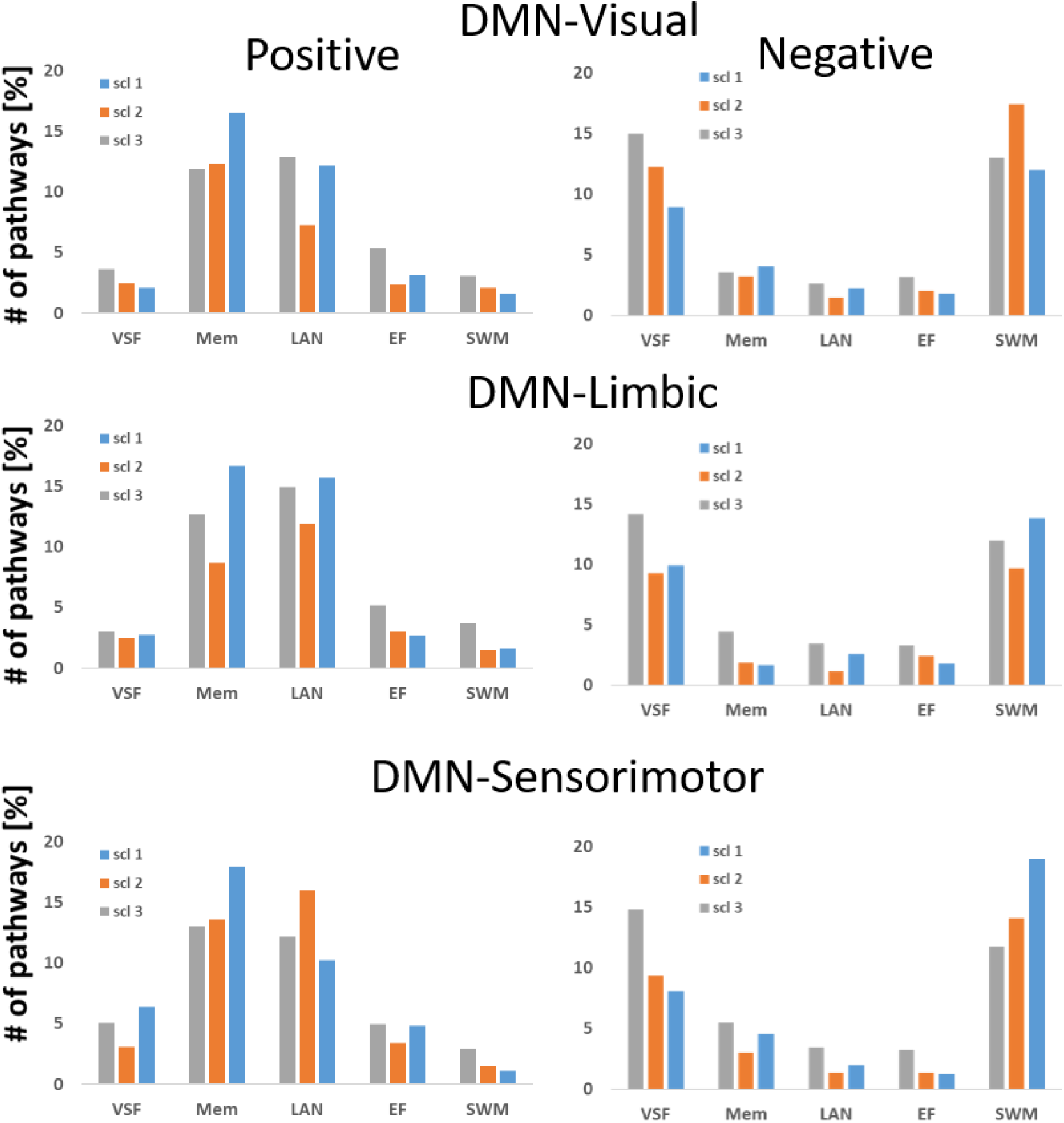
Occurrences of three-node inter-DMN pathways that correlated with five neuropsychological domains. <u>Top</u>: Pathways combining the DMN with visual regions. <u>Middle</u>: Pathways combining the DMN with limbic regions. <u>Bottom</u>: Pathways combining the DMN with sensorimotor regions. Occurrences are given in percentages of the maximum possible pathway’ number for <u>Positive</u> correlation on left and <u>Negative</u> correlation on the right. VSF - Visuospatial function; Mem-Short term memory; LAN – Language; EF-Executive function; SWM-Psychomotor speed and working memory.

**Figure 7:**
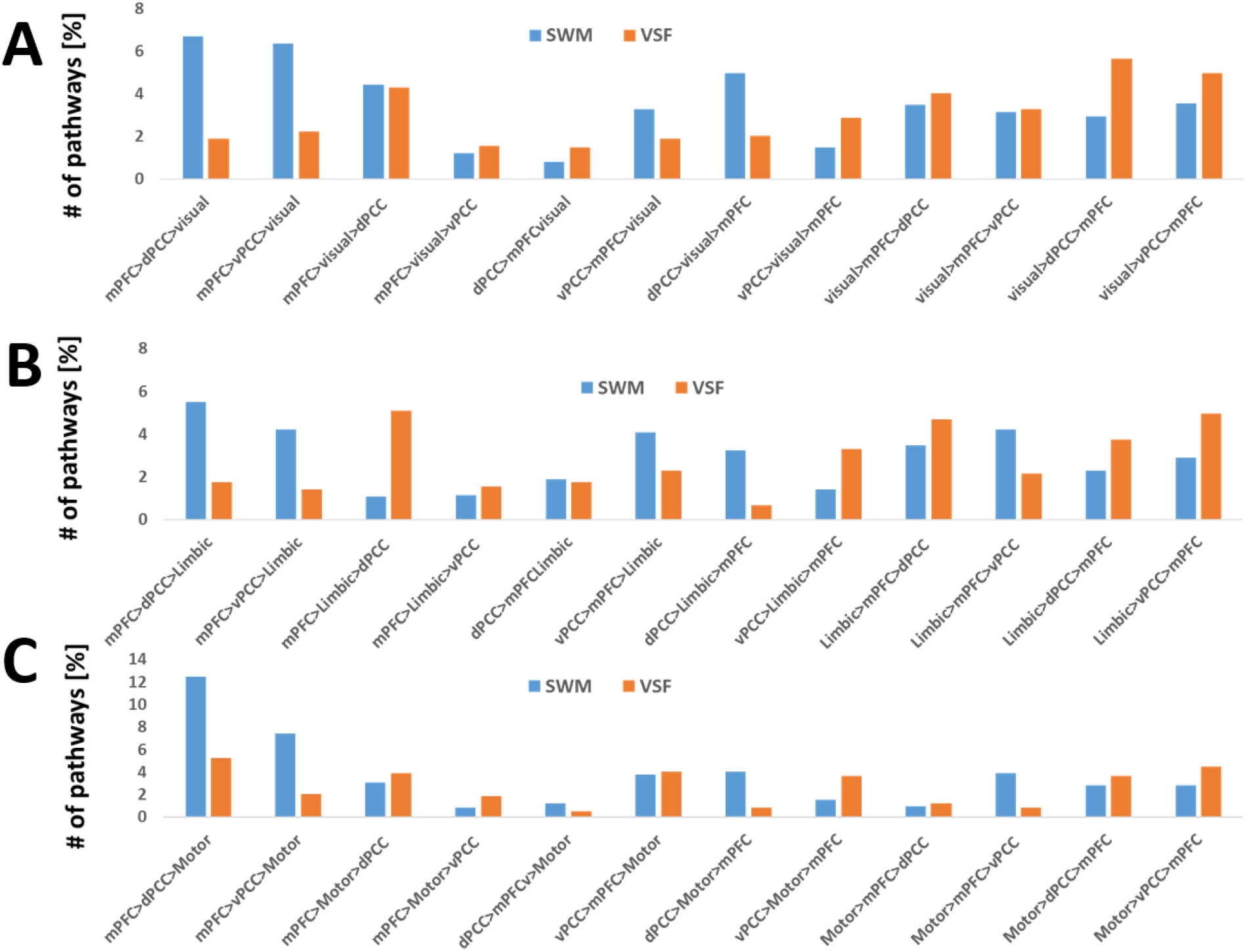
Occurrences of the three-node inter-DMN pathways that negatively correlated with psychomotor speed and working memory and with visuospatial function. Occurrences of the 12 possible pathway’ permutations are shown. A. The inter DMN-visual pathways. B. The inter DMN-Limbic pathways. C. the inter DMN-sensorimotor pathways. Note that occurrences present the sum of all three frequency scales.

**Figure 8:**
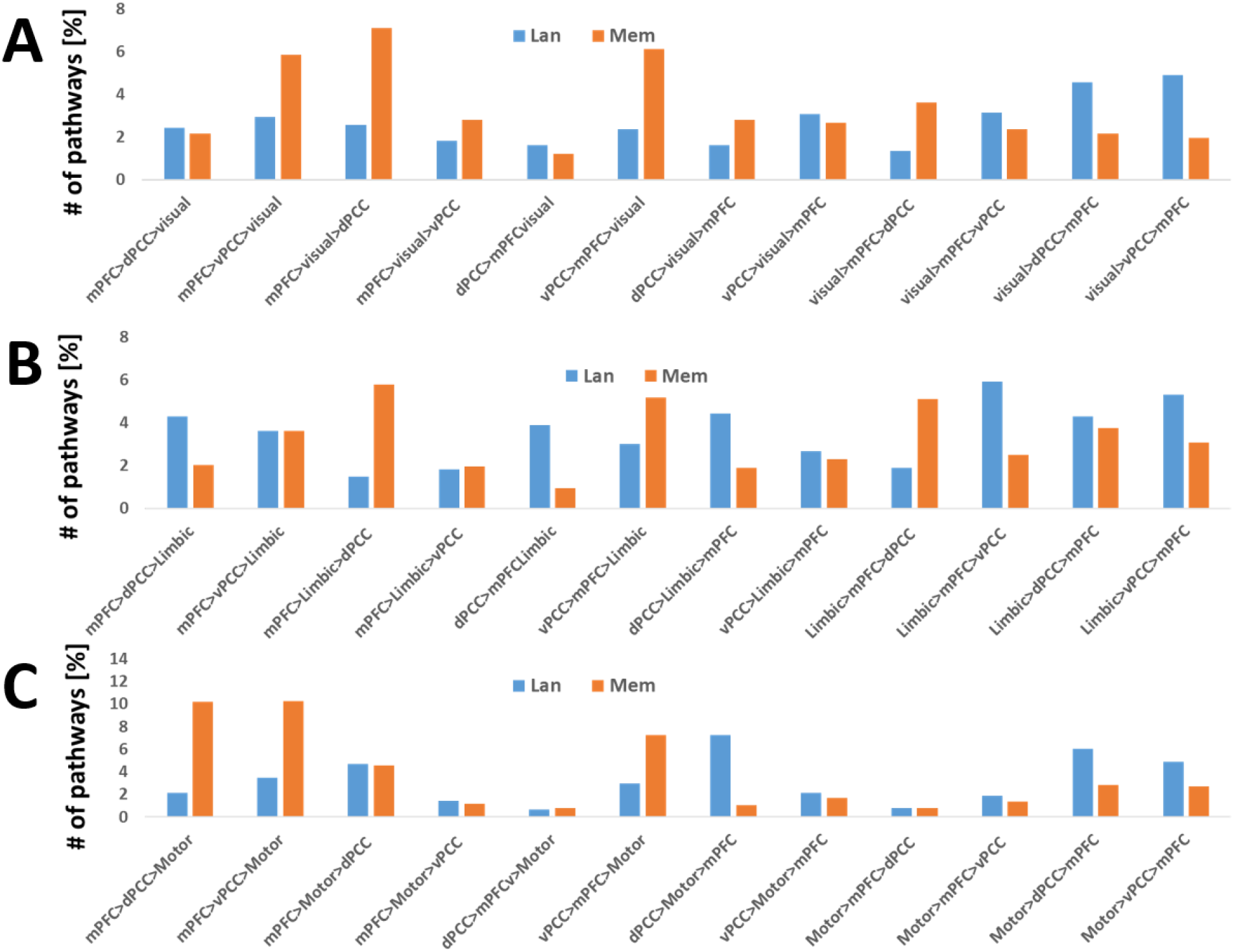
Occurrences of the three-node inter-DMN pathways that positively correlated with language and memory. Occurrences of the 12 possible pathway’ permutations are shown. A. The inter DMN-visual pathways. B. The inter DMN-Limbic pathways. C. the inter DMN-sensorimotor pathways. Note that occurrences present the sum of all three frequency scales.

## 5. Discussion

Aging is associated with dynamic changes in biological, physiological, environmental, psychological, behavioral, and social processes. The effects of aging on the brain are widespread and include among others volume shrinking, reduced dopamine and serotonin levels and vascular changes [50]. For neuroimaging macroscopic observations, we followed the line of the network degeneration hypothesis that claims that the topography of atrophy/hypometabolism follows specific brain connectivity networks [51]. We applied it to the DMN to identify degenerated or compensatory processes in elderly participants, using novel method of directed connectivity, previously developed by us [18-22]. Specifically, we aimed to develop a network model of aging that is centered in the DMN. By comparing directed pathways between healthy young and old subjects, we identified network alterations, and by calculating correlations of these pathways with a battery of neuropsychological domains, we inferred their behavioral associations.

Our main findings were the following: (i) In old versus young participants, there were fewer pathways connecting the DMN with regions in the limbic and the visual systems, and more pathways connecting the DMN with regions in the sensorimotor system. (ii) In young versus old participants, there were more pathways connecting the DMN and regions in the limbic and the visual systems that included the ventral PCC. In contrast, in old versus young participants, these pathways included the dorsal PCC. (iii) In young versus old participants, pathways between the DMN and regions within the limbic and the visual systems were at high signal frequency, while in old versus young participants they were at low frequency. (iv) Pathways connecting the sensorimotor with the DMN were sensorimotor efferent in young participants (that is lead from the sensorimotor areas towards the DMN), while in old participants they were sensorimotor afferent (that is in the opposite direction). (v) Most of the pathways that correlated with reduced speed and working memory in the old subject group were DMN efferent (lead from the DMN towards other brain locations) and (vi) Most of the pathways that correlated with increased memory were DMN efferent.

Taken together, our findings differentiated between pathways connecting regions in the sensorimotor system with the DMN and, pathways connecting regions in the visual and limbic systems with the DMN. A reduced number of pathways, a shift to the dorsal PCC, and a shift to a lower frequency in old participants characterized the pathways between the DMN and the regions in the visual and the limbic systems. In contrast, the pathways between the DMN and the sensorimotor regions were characterized by an increased number of pathways and opposite directionality between young and old participants, which also correlated with reduced psychomotor speed and working memory.

The shift from ventral to dorsal PCC in old participants is in line with previous reports that have shown different ventral and dorsal functional connectivity in amnestic, mildly cognitively impaired, and AD subjects [11]. Despite the extensive interconnections with memory, attention, and decision areas, the primary function of the PCC remains unknown. It has been suggested that the PCC controls the balance between internally and externally focused thoughts and might be a site involved in generating adaptive behavior to the changing world [9]. Specifically, due to its variation with learning, memory, reward, and task engagement, it was proposed that these modulations reflected the underlying processes of change detection and motivated subsequent shifts in behavior [52]. Furthermore, it was suggested that the ventral PCC supported internally directed thought [9], while the dorsal PCC supported externally directed thought [10]. These and other findings have led to a differentiation between the dorsal and ventral parts of the PCC and to suggestions that the dorsal PCC is implicated in dynamic control of intrinsic networks and thus in attentional focus [53]. Our findings that most of the visual regions that were found in pathways that were different between the young and old groups were within the ventral precuneus, support this account, since the ventral precuneus has been identified as a key brain region for multiple-cue judgments. This includes such features as associative memory and inference based on analysis and rules [54].

We infer that the frequency shift in pathways combining regions in the visual and the limbic systems with the DMN to a lower frequency in old participants, represents slower processing, which is in line with slower cognitive and limbic activities in old people [55]. This intuitive inference is based on our ongoing study using intracranial electroencephalography (iEEG) signals from the hippocampi of epileptic patients that show that processing times are inversely proportional to the signal frequency (paper in preparation). We speculate that a mapping between neural and BOLD frequencies exists, suggesting that BOLD signal frequencies are also related to processing times.

Most of the pathways that connect regions in the sensorimotor system with the DMN, lead from the sensorimotor areas towards the DMN in young participants and were in the opposite direction in the old participants. The later pathways were also correlated with reduced SWM. Detailed inspection of all pathways showed that the main differences between the groups were with the directionality of the mPFC. Specifically, mPFC was afferent (mPFC<) in young participants and efferent (mPFC>) in old participants. Furthermore, in most of the pathways that negatively correlated with psychomotor speed and working memory (Figure 7C), the mPFC connectivity was efferent to the sensorimotor system (e.g., mPFC>PCC>sensorimotor) indicating that these processes corresponded to reduced psychomotor speed and working memory. The mPFC is critically involved in numerous cognitive functions, including attention, inhibitory control, habit formation, working memory and long-term memory. It is however not clear what were the causes for the directionality change between young and old participants and how this change was related to reduced psychomotor speed and working memory. We note that the two findings in the old group: (i) the reduced sensorimotor efferent (sensorimotor >) and (ii) the increased sensorimotor afferent (sensorimotor <), which seem at first sight as degenerate (the former) and compensatory (the latter) processes, were not necessarily so. The fact that the sensorimotor afferent correlated with reduced psychomotor speed and working memory made these processes difficult to consider as compensatory, since we expect compensatory processes to improve and not to disprove behavior. Therefore, we speculate that these processes were uncoupled, where the reduced sensorimotor efferent may be related to reduced physical activity expected in old versus young participants, and the increased abundance of pathways in the reverse direction (sensorimotor afferent) may correspond to a higher control of the mPFC on sensorimotor activity. The former suggestion is in line with a large body of research showing the advantages of physical activity for multiple illnesses and aging [56-58], and with the evidence that the DMN is involved in these illnesses. The latter suggestion is based on findings of increased mPFC activity and motor control in the elderly [59].

## 6. Conclusions

Combining all these findings, we suggest a macroscopic model for aging that is centered in the DMN. This model may improve our understanding of aging. Further research in various population groups, including different interventions, as well as longitudinal studies monitoring changes over time, are needed to better elucidate the processes associated with aging. Meanwhile, based on the current data, we speculate that the shift of processes to lower frequencies is indicative of the reduced speed of multiple cognitive and executive processes that lead to impairments in cognitive functions [55]. The reduced sensorimotor input to the DMN, suggested as the result of reduced physical activity, and the increased need to control activity by the mPFC, causes a higher dependency on external versus internal cues. This results in a shift from ventral to dorsal PCC of inter-DMN pathways. Consequently, we speculate that one way to slow or even reverse these processes may be by increasing physical activity. Thus, this model stresses the critical importance of physical activity and suggests how it might slow aging.

## 7. Limitations

Unlike conventional analyses, coherent studies require a reference phase. Since Equations 2 and 3 were defined for a group, the reference of all the dataset (of each group separately) must be the same. It means that the group’s dataset must be acquired with the same system and that this system was not changed (e.g. upgraded) along the data collection of the group. For that reason, it is complicated to use downloaded data whose system status, are nor fully known. Such extra data could be used to validate the results and test for example intermediate age groups.

Although our data were collected at two separate sites, we assumed that the effect of site on the results was negligible. This is since: (i) Both sites had the same system, (ii) we used identical protocol and preprocessing, (iii) the null distributions for the non-parametric statistics were calculated for each group separately, thus, they included possible differences between sites, and (iv) directionality assignment was defined for each group separately.

For the multivariate pathway analysis to provide correct directed pathways, we assumed that the four regions of the pathways in each subject had comparable hemodynamic response functions (HRFs). Subjects with poor blood circulation (e.g., atherosclerotic disease) may have differential circulation, which could lead to region-specific HRF. This may have affected the results. However, all our subjects were in good health condition, therefore we do not expect regional (within subjects) differences in HRFs.

Directionality was assessed by calculated thalamocortical pathways and assuming that most pathways in the resting state are bottom-up. If this assumption is wrong, or if it is age dependent, directionality is reversed. We note that this does not affect the ventral-dorsal PCC shift or the frequency differences between the groups.

Similar to other fMRI studies, the in-scanner head motion could potentially bias the results. Major efforts were made in the acquisition and preprocessing to minimize such effects.

## Abbreviations

BOLD: Blood oxygenation level-dependent
fMRI: functional MRI

## Acknowledgements

This work was funded by the Anges Ginges Foundation, the Prusiner-Abramsky 2021 award (GG) and the Czech Ministry of Health (RJ) (grant: AZV: NV19-04-00233).

## Supplementary Material

In a separate file.

## Author Contributions

G.G.: Conception and design of the study, analysis developments, analysis, and writing of the manuscript.

R.D.: Young group data acquisition and preprocessing of all data

R.J.: Old group data acquisition and subject clinical evaluations.

O.B.: Old subject’s neuropsychological evaluations.

D.E: Writing the paper

No competing financial and/or non-financial interest for all authors

## References

[1] E. Varangis, C. G. Habeck, Q. R. Razlighi, and Y. Stern, “The Effect of Aging on Resting State Connectivity of Predefined Networks in the Brain,” (in eng), Front Aging Neurosci, vol. 11, p. 234, 2019, doi: 10.3389/fnagi.2019.00234.

[2] L. Hasher and R. T. Zacks, Working memory, comprehension,and aging: A review and a new view. (The psychology of learning and motivation). Academic Press, 1988.

[3] D. C. Park and P. Reuter-Lorenz, “The adaptive brain: aging and neurocognitive scaffolding,” (in eng), Annu Rev Psychol, vol. 60, pp. 173–96, 2009, doi: 10.1146/annurev.psych.59.103006.093656.

[4] T. A. Salthouse, “Selective review of cognitive aging,” (in eng), J Int Neuropsychol Soc, vol. 16, no. 5, pp. 754–60, Sep 2010, doi: 10.1017/S1355617710000706.

[5] J. S. Damoiseaux, “Effects of aging on functional and structural brain connectivity,” (in eng), Neuroimage, vol. 160, pp. 32–40, 10 15 2017, doi: 10.1016/j.neuroimage.2017.01.077.

[6] M. E. Raichle, “The brain’s default mode network,” Annu Rev Neurosci, vol. 38, pp. 433–47, Jul 08 2015, doi: 10.1146/annurev-neuro-071013-014030.

[7] M. D. Greicius, G. Srivastava, A. L. Reiss, and V. Menon, “Default-mode network activity distinguishes Alzheimer’s disease from healthy aging: evidence from functional MRI,” (in eng), Proc Natl Acad Sci U S A, vol. 101, no. 13, pp. 4637–42, Mar 30 2004, doi: 10.1073/pnas.0308627101.

[8] Y. I. Sheline et al., “Amyloid plaques disrupt resting state default mode network connectivity in cognitively normal elderly,” (in eng), Biol Psychiatry, vol. 67, no. 6, pp. 584–7, Mar 15 2010, doi: 10.1016/j.biopsych.2009.08.024.

[9] Y. Fan et al., “Dorsal and Ventral Posterior Cingulate Cortex Switch Network Assignment via Changes in Relative Functional Connectivity Strength to Noncanonical Networks,” (in eng), Brain Connect, vol. 9, no. 1, pp. 77–94, 02 2019, doi: 10.1089/brain.2018.0602.

[10] R. Leech, R. Braga, and D. J. Sharp, “Echoes of the brain within the posterior cingulate cortex,” (in eng), J Neurosci, vol. 32, no. 1, pp. 215–22, Jan 04 2012, doi: 10.1523/JNEUROSCI.3689-11.2012.

[11] J. Mutlu et al., “Connectivity Disruption, Atrophy, and Hypometabolism within Posterior Cingulate Networks in Alzheimer’s Disease,” (in eng), Front Neurosci, vol. 10, p. 582, 2016, doi: 10.3389/fnins.2016.00582.

[12] J. R. Andrews-Hanna et al., “Disruption of large-scale brain systems in advanced aging,” (in eng), Neuron, vol. 56, no. 5, pp. 924–35, Dec 06 2007, doi: 10.1016/j.neuron.2007.10.038.

[13] A. M. Staffaroni et al., “The Longitudinal Trajectory of Default Mode Network Connectivity in Healthy Older Adults Varies As a Function of Age and Is Associated with Changes in Episodic Memory and Processing Speed,” (in eng), J Neurosci, vol. 38, no. 11, pp. 2809–2817, 03 2018, doi: 10.1523/JNEUROSCI.3067-17.2018.

[14] A. C. Simioni, A. Dagher, and L. K. Fellows, “Compensatory striatal-cerebellar connectivity in mild-moderate Parkinson’s disease,” (in eng), Neuroimage Clin, vol. 10, pp. 54–62, 2016, doi: 10.1016/j.nicl.2015.11.005.

[15] F. G. Hillary et al., “The rich get richer: brain injury elicits hyperconnectivity in core subnetworks,” (in eng), PLoS One, vol. 9, no. 8, p. e104021, 2014, doi: 10.1371/journal.pone.0104021.

[16] C. Yang, S. Zhong, X. Zhou, L. Wei, L. Wang, and S. Nie, “The Abnormality of Topological Asymmetry between Hemispheric Brain White Matter Networks in Alzheimer’s Disease and Mild Cognitive Impairment,” Front Aging Neurosci, vol. 9, p. 261, 2017, doi: 10.3389/fnagi.2017.00261.

[17] R. A. Menke et al., “Increased functional connectivity common to symptomatic amyotrophic lateral sclerosis and those at genetic risk,” (in eng), J Neurol Neurosurg Psychiatry, vol. 87, no. 6, pp. 580–8, 06 2016, doi: 10.1136/jnnp-2015-311945.

[18] G. Goelman and R. Dan, “Multiple-region directed functional connectivity based on phase delays,” Hum Brain Mapp, vol. 38, no. 3, pp. 1374–1386, Mar 2017, doi: 10.1002/hbm.23460.

[19] G. Goelman, R. Dan, and T. Keadan, “Characterizing directed functional pathways in the visual system by multivariate nonlinear coherence of fMRI data,” Sci Rep, vol. 8, no. 1, p. 16362, Nov 5 2018, doi: 10.1038/s41598-018-34672-5.

[20] G. Goelman, R. Dan, G. Stossel, H. Tost, A. Meyer-Lindenberg, and E. Bilek, “Bidirectional signal exchanges and their mechanisms during joint attention interaction - A hyperscanning fMRI study,” Neuroimage, vol. 198, pp. 242–254, Sep 2019, doi: 10.1016/j.neuroimage.2019.05.028.

[21] G. Goelman, R. Dan, F. Růžička, O. Bezdicek, and R. Jech, “Altered sensorimotor fMRI directed connectivity in Parkinson’s disease patients,” (in eng), Eur J Neurosci, vol. 53, no. 6, pp. 1976–1987, 03 2021, doi: 10.1111/ejn.15053.

[22] G. Goelman, R. Dan, F. Růžička, O. Bezdicek, and R. Jech, “Asymmetry of the insula-sensorimotor circuit in Parkinson’s disease,” (in eng), Eur J Neurosci, Aug 27 2021, doi: 10.1111/ejn.15432.

[23] R. Dan et al., “Sex differences during emotion processing are dependent on the menstrual cycle phase,” Psychoneuroendocrinology, vol. 100, pp. 85–95, Feb 2019, doi: 10.1016/j.psyneuen.2018.09.032.

[24] R. Dan et al., “Impact of dopamine and cognitive impairment on neural reactivity to facial emotion in Parkinson’s disease,” Eur Neuropsychopharmacol, vol. 29, no. 11, pp. 1258–1272, Nov 2019, doi: 10.1016/j.euroneuro.2019.09.003.

[25] O. Bezdicek et al., “Czech version of the Trail Making Test: normative data and clinical utility,” (in eng), Arch Clin Neuropsychol, vol. 27, no. 8, pp. 906–14, Dec 2012, doi: 10.1093/arclin/acs084.

[26] D. Wechsler, Wechsler Adult Intelligence Scale-3rd ed. (WAIS-3). San Antonio, TX: Harcourt Assess., 1997.

[27] J. Michalec et al., “Standardization of the Czech version of the Tower of London test - Administration, scoring, validity,” Ces. a Slov. Neurol. a Neurochir, no. 77, pp. 596–601, 2014.

[28] T. Nikolai et al., “Tests of Verbal Fluency, Czech Normative Study in Older Patients,” Česká a Slov. Neurol. a Neurochir., no. 78, 2015.

[29] N. Zemanová et al., “Validity Study of the Boston Naming Test Czech Version,” Česká a Slov. Neurol. a Neurochir, vol. 79, pp. 307–316, 2016.

[30] O. Bezdicek, A. M. Rosická, J. Mana, D. J. Libon, M. Kopeček, and H. Georgi, “The 30-item and 15-item Boston naming test Czech version: Item response analysis and normative values for healthy older adults,” (in eng), J Clin Exp Neuropsychol, vol. 43, no. 9, pp. 890–905, 11 2021, doi: 10.1080/13803395.2022.2029360.

[31] O. Bezdicek et al., “Czech version of Rey Auditory Verbal Learning test: normative data,” (in eng), Neuropsychol Dev Cogn B Aging Neuropsychol Cogn, vol. 21, no. 6, pp. 693–721, 2014, doi: 10.1080/13825585.2013.865699.

[32] R. H. Benedict, D. Schretlen, L. Groninger, M. Dobraski S, and B. hpritz, “Revision of the Brief Visuospatial Memory Test: Studies of normal performance, reliability, and validity,” Psychol. Assess., vol. 8, pp. 145–153, 1996.

[33] F. Havlík et al., “Brief Visuospatial Memory Test-Revised: normative data and clinical utility of learning indices in Parkinson’s disease,” (in eng), J Clin Exp Neuropsychol, vol. 42, no. 10, pp. 1099–1110, 12 2020, doi: 10.1080/13803395.2020.1845303.

[34] D. R. Royall, J. A. Cordes, and M. Polk, “CLOX: an executive clock drawing task,” (in eng), J Neurol Neurosurg Psychiatry, vol. 64, no. 5, pp. 588–94, May 1998, doi: 10.1136/jnnp.64.5.588.

[35] J. L. Woodard, R. H. Benedict, T. A. Salthouse, J. P. Toth, D. J. Zgaljardic, and H. E. Hancock, “Normative data for equivalent, parallel forms of the Judgment of Line Orientation Test,” (in eng), J Clin Exp Neuropsychol, vol. 20, no. 4, pp. 457–62, Aug 1998, doi: 10.1076/jcen.20.4.457.1473.

[36] S. R. Solomon and S. S. Sawilowsky, “Impact of Rank-Based Normalizing Transformations on the Accuracy of Test Scores,” J. Mod. Appl. Stat. Methods, vol. 8, pp. 448–462, 2009.

[37] S. Whitfield-Gabrieli and A. Nieto-Castanon, “Conn: a functional connectivity toolbox for correlated and anticorrelated brain networks,” Brain Connect, vol. 2, no. 3, pp. 125–41, 2012, doi: 10.1089/brain.2012.0073.

[38] Y. Behzadi, K. Restom, J. Liau, and T. T. Liu, “A component based noise correction method (CompCor) for BOLD and perfusion based fMRI,” (in eng), Neuroimage, vol. 37, no. 1, pp. 90–101, Aug 2007, doi: 10.1016/j.neuroimage.2007.04.042.

[39] J. D. Power, A. Mitra, T. O. Laumann, A. Z. Snyder, B. L. Schlaggar, and S. E. Petersen, “Methods to detect, characterize, and remove motion artifact in resting state fMRI,” Neuroimage, vol. 84, pp. 320–41, Jan 1 2014, doi: 10.1016/j.neuroimage.2013.08.048.

[40] K. Murphy, R. M. Birn, D. A. Handwerker, T. B. Jones, and P. A. Bandettini, “The impact of global signal regression on resting state correlations: are anti-correlated networks introduced?,” (in eng), Neuroimage, vol. 44, no. 3, pp. 893–905, Feb 01 2009, doi: 10.1016/j.neuroimage.2008.09.036.

[41] X. J. Chai, A. N. Castañón, D. Ongür, and S. Whitfield-Gabrieli, “Anticorrelations in resting state networks without global signal regression,” (in eng), Neuroimage, vol. 59, no. 2, pp. 1420–8, Jan 16 2012, doi: 10.1016/j.neuroimage.2011.08.048.

[42] C. J. Stam, G. Nolte, and A. Daffertshofer, “Phase lag index: assessment of functional connectivity from multi channel EEG and MEG with diminished bias from common sources,” Human brain mapping, vol. 28, no. 11, pp. 1178–93, Nov 2007, doi: 10.1002/hbm.20346.

[43] C. J. Stam and E. C. van Straaten, “Go with the flow: use of a directed phase lag index (dPLI) to characterize patterns of phase relations in a large-scale model of brain dynamics,” NeuroImage, vol. 62, no. 3, pp. 1415–28, Sep 2012, doi: 10.1016/j.neuroimage.2012.05.050.

[44] C. Torrence and G. P. Compo, “Wavelet Analysis,” Bull. Amer. Meteor. Soc., vol. 79, pp. 61–78, 1998.

[45] C. Torrence and P. Webster, “Interdecadal changes in the ENSO-Monsoon system,” J Clim, vol. 12, pp. 2679–2690, 1999.

[46] K. Muller, G. Lohmann, J. Neumann, M. Grigutsch, T. Mildner, and D. Y. von Cramon, “Investigating the wavelet coherence phase of the BOLD signal,” J Magn Reson Imaging, vol. 20, no. 1, pp. 145–52, Jul 2004, doi: 10.1002/jmri.20064.

[47] C. Torrence and P. Webster, “Interdecadal changes in the ENSO-Monsoon system,” J Clim, vol. 12, pp. 2679–2690, 1999.

[48] M. Joliot et al., “AICHA: An atlas of intrinsic connectivity of homotopic areas,” Journal of neuroscience methods, vol. 254, pp. 46–59, Oct 30 2015, doi: 10.1016/j.jneumeth.2015.07.013.

[49] N. Tzourio-Mazoyer et al., “Automated anatomical labeling of activations in SPM using a macroscopic anatomical parcellation of the MNI MRI single-subject brain,” NeuroImage, vol. 15, no. 1, pp. 273–89, Jan 2002, doi: 10.1006/nimg.2001.0978.

[50] R. Peters, “Ageing and the brain,” (in eng), Postgrad Med J, vol. 82, no. 964, pp. 84–8, Feb 2006, doi: 10.1136/pgmj.2005.036665.

[51] W. W. Seeley, R. K. Crawford, J. Zhou, B. L. Miller, and M. D. Greicius, “Neurodegenerative diseases target large-scale human brain networks,” (in eng), Neuron, vol. 62, no. 1, pp. 42–52, Apr 16 2009, doi: 10.1016/j.neuron.2009.03.024.

[52] J. M. Pearson, S. R. Heilbronner, D. L. Barack, B. Y. Hayden, and M. L. Platt, “Posterior cingulate cortex: adapting behavior to a changing world,” (in eng), Trends Cogn Sci, vol. 15, no. 4, pp. 143–51, Apr 2011, doi: 10.1016/j.tics.2011.02.002.

[53] R. Leech and D. J. Sharp, “The role of the posterior cingulate cortex in cognition and disease,” (in eng), Brain, vol. 137, no. Pt 1, pp. 12–32, Jan 2014, doi: 10.1093/brain/awt162.

[54] L. K. Wirebring, S. Stillesjö, J. Eriksson, P. Juslin, and L. Nyberg, “A Similarity-Based Process for Human Judgment in the Parietal Cortex,” (in eng), Front Hum Neurosci, vol. 12, p. 481, 2018, doi: 10.3389/fnhum.2018.00481.

[55] T. A. Salthouse, “The processing-speed theory of adult age differences in cognition,” (in eng), Psychol Rev, vol. 103, no. 3, pp. 403–28, Jul 1996, doi: 10.1037/0033-295x.103.3.403.

[56] A. Rebelo-Marques et al., “Aging Hallmarks: The Benefits of Physical Exercise,” (in eng), Front Endocrinol (Lausanne), vol. 9, p. 258, 2018, doi: 10.3389/fendo.2018.00258.

[57] D. Tavoian, D. W. Russ, L. A. Consitt, and B. C. Clark, “Perspective: Pragmatic Exercise Recommendations for Older Adults: The Case for Emphasizing Resistance Training,” (in eng), Front Physiol, vol. 11, p. 799, 2020, doi: 10.3389/fphys.2020.00799.

[58] M. Izquierdo et al., “International Exercise Recommendations in Older Adults (ICFSR): Expert Consensus Guidelines,” (in eng), J Nutr Health Aging, vol. 25, no. 7, pp. 824–853, 2021, doi: 10.1007/s12603-021-1665-8.

[59] L. Zapparoli, M. Mariano, and E. Paulesu, “How the motor system copes with aging: a quantitative meta-analysis of the effect of aging on motor function control,” (in eng), Commun Biol, vol. 5, no. 1, p. 79, 01 20 2022, doi: 10.1038/s42003-022-03027-2.

