## Supplement Table and Figures for "Older age is associated with a shift from ventral to dorsal PCC in pathways connecting the DMN with visual and limbic areas and a directionality shift in pathways connecting the DMN with sensorimotor areas"

**Table 1**

| **PW number** | **Pathway** |
| --- | --- |
| 1 | R1-R2-R3-X |
| 2 | R2-R3-X-R1 |
| 3 | R3-X-R1-R2 |
| 4 | X-R1-R2-R3 |
| 5 | R1-R2-X-R3 |
| 6 | R2-X-R3-R1 |
| 7 | R3-R1-R2-X |
| 8 | X-R3-R1-R2 |
| 9 | R1-R3-X-R2 |
| 10 | R3-X-R2-R1 |
| 11 | X-R2-R1-R3 |
| 12 | R2-R1-R3-X |
| 13 | R1-X-R3-R2 |
| 14 | R2-R1-X-R3 |
| 15 | R3-R2-R1-X |
| 16 | X-R3-R2-R1 |
| 17 | R1-R3-R2-X |
| 18 | R2-X-R1-R3 |
| 19 | R3-R2-X-R1 |
| 20 | X-R1-R3-R2 |
| 21 | R1-X-R2-R3 |
| 22 | R2-R3-R1-X |
| 23 | R3-R1-X-R2 |
| 24 | X-R2-R3-R1 |

**Table 1:** A list of all four-node pathways. R1 to R3 are three predefined regions, while X is a one of the 384 AICHA (Atlas of Intrinsic Connectivity of Homotopic Areas) regions.

| **Young** | Visual | | Limbic | | Motor | |
| --- | --- | --- | --- | --- | --- | --- |
|  | Scale 1 | Scale 2 | Scale 1 | Scale 2 | Scale 1 | Scale 2 |
| mPFC>dPCC>system |  |  |  |  |  |  |
| mPFC>vPCC>system | 9.65 | 13.94 |  | 2.55 |  | 1.74 |
| mPFC>system>dPCC |  | 1.88 |  |  |  | 1.47 |
| mPFC>system>vPCC | 3.35 | 14.61 |  | 2.82 |  | 1.88 |
| dPCC>mPFC>system | 2.41 |  | 1.07 | 1.07 |  |  |
| vPCC>mPFC>system |  |  |  |  |  |  |
| dPCC>system>mPFC | 1.21 |  |  |  | 2.95 | 1.21 |
| vPCC>system>mPFC | 1.34 | 1.47 |  |  |  |  |
| system>mPFC>dPCC |  |  |  |  |  |  |
| system>mPFC>vPCC |  | 4.83 |  | 4.42 | 1.34 | 2.41 |
| system>dPCC>mPFC |  |  | 1.07 |  | 4.83 | 1.34 |
| system>vPCC>mPFC | 2.28 |  |  |  | 2.41 |  |

| **Old** | Visual | | Limbic | | Motor | |
| --- | --- | --- | --- | --- | --- | --- |
|  | Scale 1 | Scale 2 | Scale 1 | Scale 2 | Scale 1 | Scale 2 |
| mPFC>dPCC>system |  |  |  |  | 10.46 | 5.23 |
| mPFC>vPCC>system | 4.16 | 1.47 | 1.74 |  | 3.89 | 1.74 |
| mPFC>system>dPCC | 2.28 | 1.07 | 1.47 |  | 1.34 | 2.01 |
| mPFC>system>vPCC |  | 1.21 |  |  |  |  |
| dPCC>mPFC>system |  |  |  |  |  |  |
| vPCC>mPFC>system | 1.61 |  | 1.07 |  | 8.04 |  |
| dPCC>system>mPFC |  |  |  |  |  | 2.01 |
| vPCC>system>mPFC |  |  |  |  |  |  |
| system>mPFC>dPCC | 2.01 |  |  |  |  |  |
| system>mPFC>vPCC | 1.07 |  | 1.61 |  | 1.07 |  |
| system>dPCC>mPFC |  |  |  |  |  |  |
| system>vPCC>mPFC |  |  |  |  |  |  |

**Table 2:** Percentages of a 3-node one-group DMN↔system (visual, limbic and sensorimotor). A. The young participant group. B. The old participant group. Cells with pathways of 1% of the maximum number of possible pathways are more are shown.

A. Inter DMN-Limbic Correlations

| **DMN-Limbic**  **Positive Correlations** | SWM | | | EF | | | Lan | | | Mem | | | VSF | | |
| --- | --- | --- | --- | --- | --- | --- | --- | --- | --- | --- | --- | --- | --- | --- | --- |
| Scale | 1 | 2 | 3 | 1 | 2 | 3 | 1 | 2 | 3 | 1 | 2 | 3 | 1 | 2 | 3 |
| mPFC>dPCC>system |  |  |  |  |  |  | 1.2 | 2.0 | 1.1 |  |  |  | 1.3 |  |  |
| mPFC>vPCC>system |  |  |  |  |  |  | 1.5 | 1.2 |  |  | 1.3 | 1.5 |  |  |  |
| mPFC>system>dPCC |  |  |  |  | 1.1 |  |  |  |  | 3.9 |  | 1.1 |  |  |  |
| mPFC>system>vPCC |  |  |  |  |  |  |  |  | 1.0 |  |  |  |  |  |  |
| dPCC>mPFC>system |  |  |  |  |  |  | 1.6 | 1.3 |  |  |  |  |  |  |  |
| vPCC>mPFC>system |  |  |  |  |  |  | 1.3 | 1.4 |  | 2.1 | 1.7 | 1.3 |  |  |  |
| dPCC>system>mPFC |  |  |  |  |  |  | 1.6 | 1.3 | 1.5 |  |  |  |  |  |  |
| vPCC>system>mPFC |  |  |  |  |  |  |  | 1.2 |  | 1.2 |  |  |  |  |  |
| system>mPFC>dPCC |  |  |  |  |  |  | 1.2 |  |  | 3.9 |  |  |  |  |  |
| system>mPFC>vPCC |  |  |  |  |  |  | 4.3 |  | 1.0 |  |  |  |  |  |  |
| system>dPCC>mPFC |  |  | 1.7 |  |  |  |  |  | 3.2 |  |  | 3.4 |  |  |  |
| system>vPCC>mPFC |  |  |  |  |  | 1.4 | 0.9 | 1.4 | 2.9 |  |  | 1.1 |  |  |  |

| **DMN-Limbic**  **Negative Correlations** | SWM | | | EF | | | Lan | | | Mem | | | VSF | | |
| --- | --- | --- | --- | --- | --- | --- | --- | --- | --- | --- | --- | --- | --- | --- | --- |
| Scale | 1 | 2 | 3 | 1 | 2 | 3 | 1 | 2 | 3 | 1 | 2 | 3 | 1 | 2 | 3 |
| mPFC>dPCC>system | 4.0 |  |  |  |  |  |  |  |  |  |  |  |  |  |  |
| mPFC>vPCC>system | 1.7 |  | 1.8 |  |  |  |  |  |  |  |  |  |  |  |  |
| mPFC>system>dPCC |  |  |  |  |  |  |  |  |  |  |  |  | 2.1 | 2.0 |  |
| mPFC>system>vPCC |  |  |  |  |  |  |  |  |  |  |  |  |  |  |  |
| dPCC>mPFC>system |  |  |  |  |  |  |  |  |  |  |  |  |  | 1.3 |  |
| vPCC>mPFC>system | 1.4 | 1.0 | 1.7 |  |  |  |  |  |  |  |  |  |  | 1.3 |  |
| dPCC>system>mPFC |  | 2.4 |  |  |  |  |  |  |  |  |  |  |  |  |  |
| vPCC>system>mPFC |  |  |  |  |  |  |  |  |  |  |  |  | 1.5 |  | 1.2 |
| system>mPFC>dPCC | 2.0 |  | 1.3 |  |  |  |  |  |  |  |  |  | 3.1 |  | 1.5 |
| system>mPFC>vPCC | 2.6 | 1.2 |  |  |  |  |  |  |  |  |  |  |  |  | 1.1 |
| system>dPCC>mPFC |  |  | 2.0 |  |  |  |  |  | 1.1 |  |  | 1.1 |  |  | 3.1 |
| system>vPCC>mPFC |  | 1.1 | 1.3 |  |  | 1.5 |  |  |  |  |  | 1.7 |  | 1.3 | 3.4 |

B. Inter DMN-Visual Correlations

| **DMN-visual**  **Positive Correlations** | SWM | | | EF | | | Lan | | | Mem | | | VSF | | |
| --- | --- | --- | --- | --- | --- | --- | --- | --- | --- | --- | --- | --- | --- | --- | --- |
| Scale | 1 | 2 | 3 | 1 | 2 | 3 | 1 | 2 | 3 | 1 | 2 | 3 | 1 | 2 | 3 |
| mPFC>dPCC>system |  |  |  |  |  |  |  | 1.2 |  |  |  |  |  |  |  |
| mPFC>vPCC>system |  |  |  |  |  |  | 1.7 |  |  | 2.3 | 1.6 | 1.9 |  |  |  |
| mPFC>system>dPCC |  |  |  |  |  |  |  |  |  | 2.8 | 2.7 | 1.6 |  |  |  |
| mPFC>system>vPCC |  |  |  |  |  |  | 1.3 |  |  |  |  | 1.4 |  |  |  |
| dPCC>mPFC>system |  |  |  |  |  |  | 1.2 |  |  |  |  |  |  |  |  |
| vPCC>mPFC>system |  |  |  |  |  |  |  | 1.0 |  | 2.9 | 2.0 | 1.1 |  |  |  |
| dPCC>system>mPFC |  |  |  |  |  |  | 1.1 |  |  | 2.4 |  |  |  |  |  |
| vPCC>system>mPFC |  |  |  |  |  |  |  | 0.7 | 1.7 | 1.8 |  |  |  |  |  |
| system>mPFC>dPCC |  |  |  |  |  |  |  |  |  | 1.2 | 2.3 |  |  |  |  |
| system>mPFC>vPCC |  |  |  |  |  |  | 1.7 |  |  |  | 1.0 | 1.0 |  |  |  |
| system>dPCC>mPFC |  |  |  |  |  |  | 1.5 |  | 2.5 |  |  | 1.1 |  |  |  |
| system>vPCC>mPFC |  |  |  |  |  | 1.4 |  | 1.2 | 3.5 |  |  | 1.3 |  |  | 1.1 |

| **DMN-Visual**  **Negative Correlations** | SWM | | | EF | | | Lan | | | Mem | | | VSF | | |
| --- | --- | --- | --- | --- | --- | --- | --- | --- | --- | --- | --- | --- | --- | --- | --- |
| Scale | 1 | 2 | 3 | 1 | 2 | 3 | 1 | 2 | 3 | 1 | 2 | 3 | 1 | 2 | 3 |
| mPFC>dPCC>system | 1.5 | 4.0 | 1.2 |  |  |  |  |  |  |  |  |  | 1.2 |  |  |
| mPFC>vPCC>system | 1.4 | 2.5 | 2.5 |  |  |  |  |  |  |  |  |  | 1.1 |  |  |
| mPFC>system>dPCC | 3.4 |  |  |  |  |  |  |  |  |  |  |  | 1.3 | 1.3 | 1.6 |
| mPFC>system>vPCC |  |  |  |  |  |  |  |  |  |  |  |  |  |  |  |
| dPCC>mPFC>system |  |  |  |  |  |  |  |  |  |  |  |  |  |  |  |
| vPCC>mPFC>system | 1.6 |  |  |  |  |  |  |  |  |  |  |  |  | 1.0 |  |
| dPCC>system>mPFC |  | 4.0 |  |  |  |  |  |  |  |  |  |  | 1.2 |  |  |
| vPCC>system>mPFC |  |  |  |  |  |  |  |  |  |  |  |  | 1.5 |  |  |
| system>mPFC>dPCC | 2.3 |  | 1.2 |  |  |  |  |  |  |  |  |  |  |  | 2.5 |
| system>mPFC>vPCC |  | 1.8 |  |  |  |  |  |  |  |  |  |  |  | 2.5 |  |
| system>dPCC>mPFC |  |  | 2.1 |  |  |  |  |  |  |  |  |  |  | 2.1 | 3.4 |
| system>vPCC>mPFC |  | 1.3 | 1.8 |  |  |  |  |  |  |  |  | 1.9 |  | 1.5 | 3.4 |

C. Inter DMN-Sensorimotor Correlations

| **DMN-SM**  **Positive Correlations** | SWM | | | EF | | | Lan | | | Mem | | | VSF | | |
| --- | --- | --- | --- | --- | --- | --- | --- | --- | --- | --- | --- | --- | --- | --- | --- |
| Scale | 1 | 2 | 3 | 1 | 2 | 3 | 1 | 2 | 3 | 1 | 2 | 3 | 1 | 2 | 3 |
| mPFC>dPCC>system |  |  |  | 2.0 | 1.7 |  |  | 1.1 |  | 7.0 | 2.1 | 1.1 | 4 |  |  |
| mPFC>vPCC>system |  |  |  |  |  |  | 1.6 |  | 1.1 | 2.0 | 5.8 | 2.4 |  |  |  |
| mPFC>system>dPCC |  |  |  |  |  |  |  | 2.3 | 1.7 |  | 2.3 | 1.5 |  |  |  |
| mPFC>system>vPCC |  |  |  |  |  |  |  |  |  |  |  |  |  |  |  |
| dPCC>mPFC>system |  |  |  |  |  |  |  |  |  |  |  |  |  |  |  |
| vPCC>mPFC>system |  |  |  |  |  |  | 1.3 |  |  | 4.3 |  |  |  |  |  |
| dPCC>system>mPFC |  |  |  |  |  |  | 1.7 | 5.0 |  |  |  |  |  |  |  |
| vPCC>system>mPFC |  |  |  |  |  |  |  | 1.0 |  | 1.2 |  |  |  |  |  |
| system>mPFC>dPCC |  |  |  |  |  |  |  |  |  |  |  |  |  |  |  |
| system>mPFC>vPCC |  |  |  |  |  |  |  | 1.0 |  |  |  |  |  |  |  |
| system>dPCC>mPFC |  |  |  |  |  | 1.5 | 2.1 |  | 2.9 |  |  | 1.9 |  |  |  |
| system>vPCC>mPFC |  |  |  |  |  |  |  | 2.7 | 1.9 |  |  | 2.1 |  |  |  |

| **DMN-SM**  **Negative Correlations** | SWM | | | EF | | | Lan | | | Mem | | | VSF | | |
| --- | --- | --- | --- | --- | --- | --- | --- | --- | --- | --- | --- | --- | --- | --- | --- |
| Scale | 1 | 2 | 3 | 1 | 2 | 3 | 1 | 2 | 3 | 1 | 2 | 3 | 1 | 2 | 3 |
| mPFC>dPCC>system | 8.7 | 2.7 | 1.1 |  |  |  |  |  |  |  |  |  | 2.3 | 1.3 | 1.6 |
| mPFC>vPCC>system | 3.8 | 1.7 | 1.9 |  |  |  |  |  |  |  |  |  |  |  | 1.1 |
| mPFC>system>dPCC | 1.7 |  |  |  |  |  |  |  |  |  |  |  | 1.2 | 2.3 |  |
| mPFC>system>vPCC |  |  |  |  |  |  |  |  |  |  |  |  |  |  |  |
| dPCC>mPFC>system |  |  |  |  |  |  |  |  |  |  |  |  |  |  |  |
| vPCC>mPFC>system | 2.3 | 1.2 |  |  |  |  |  |  |  |  |  |  | 1.5 |  | 1.7 |
| dPCC>system>mPFC |  | 2.8 | 1.2 |  |  |  |  |  |  |  |  |  |  |  |  |
| vPCC>system>mPFC |  |  |  |  |  |  |  |  |  |  |  |  | 1.1 |  | 1.6 |
| system>mPFC>dPCC |  |  |  |  |  |  |  |  |  |  |  |  |  |  | 1.1 |
| system>mPFC>vPCC |  | 1.1 | 2.3 |  |  |  |  |  |  |  |  |  |  |  |  |
| system>dPCC>mPFC |  |  | 2.1 |  |  |  |  |  |  |  |  |  |  |  | 2.7 |
| system>vPCC>mPFC |  | 1.3 | 1.1 |  |  |  |  |  |  |  |  | 1.7 |  | 1.0 | 3.2 |

**Table 3:** Percentages of a 3-node DMN↔system pathways of the old subject group that correlated with neurophysiological categories. Cells with pathways of >1% of the maximum number of possible pathways, are shown.

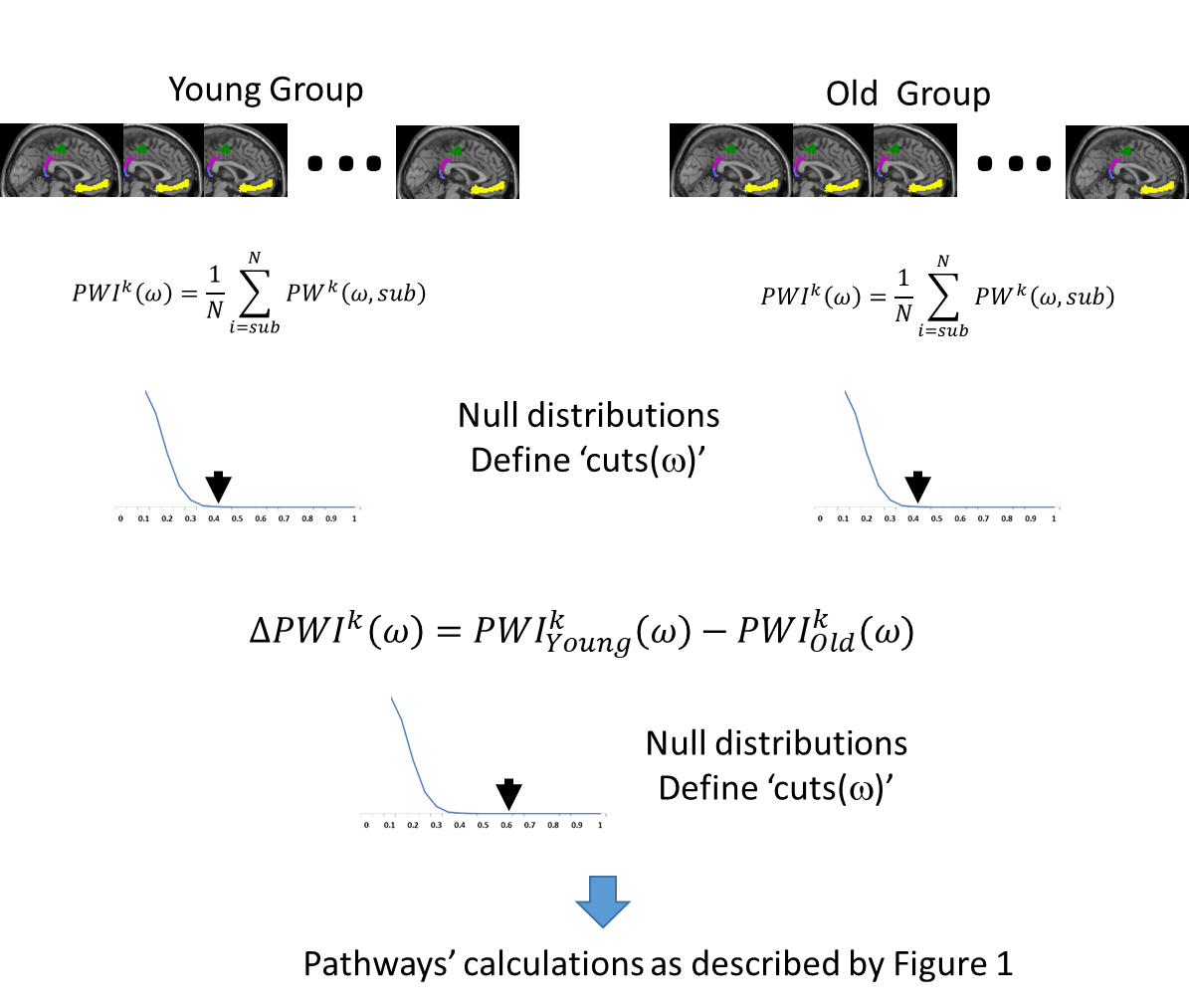

Figure 1: Illustration of the stages used to obtain group and between groups significant pathways.

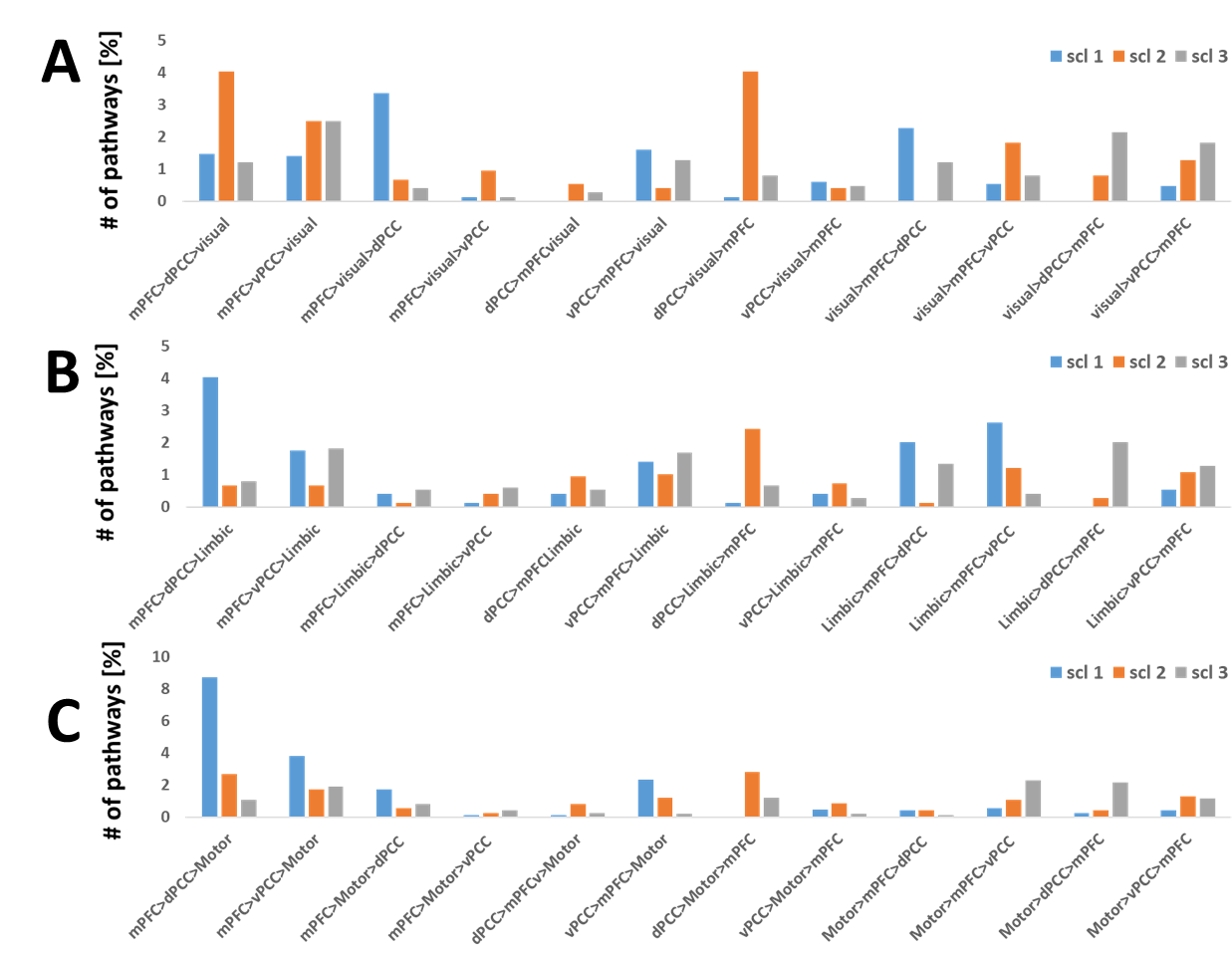

Figure 2: Occurrences of the three-node inter-DMN pathways that negatively correlated with speed and working memory. Occurrences of the 12 possible pathway' permutations are shown. A. The inter DMN-visual pathways. B. The inter DMN-Limbic pathways. C. The inter DMN-sensorimotor pathways.

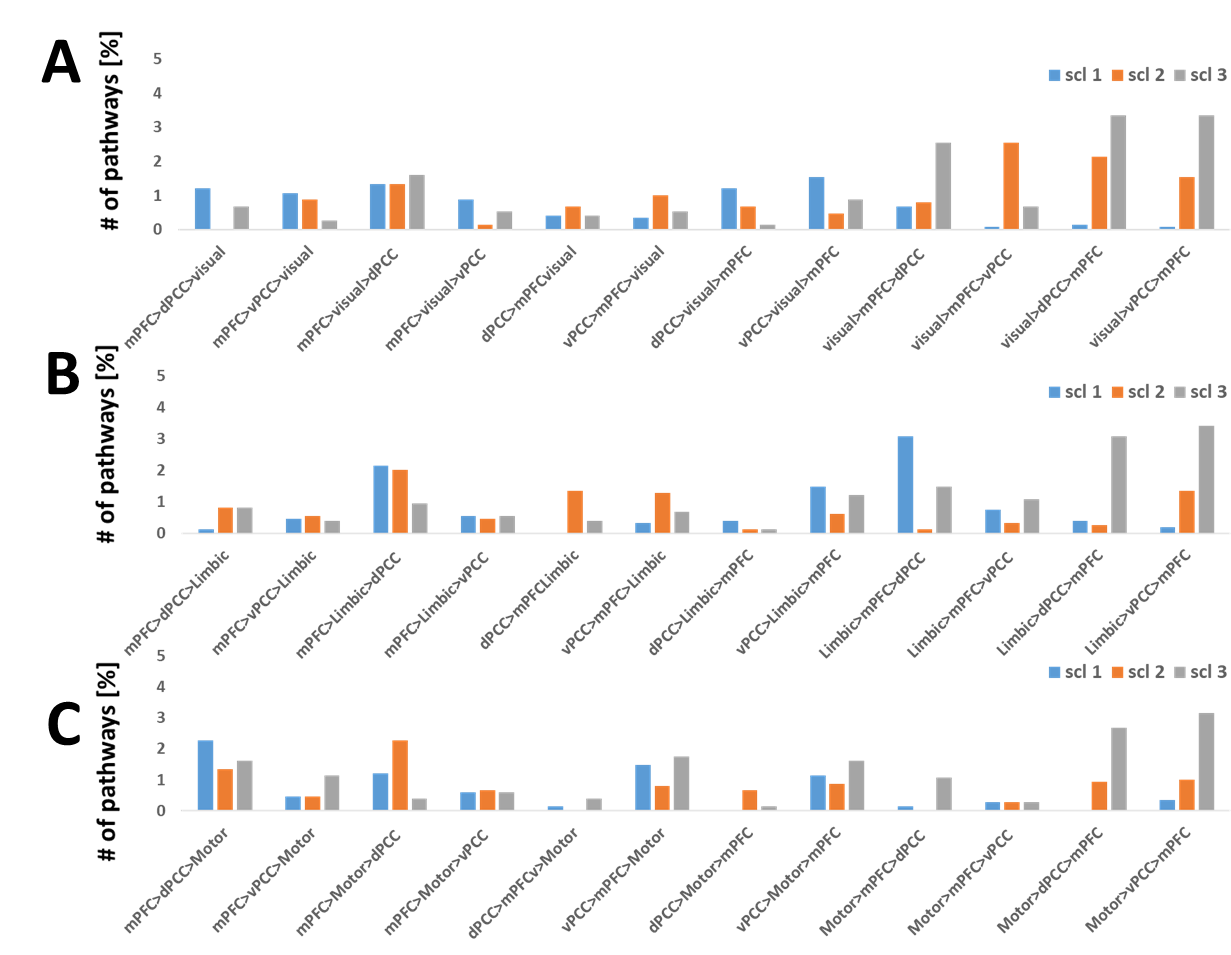

Figure 3: Occurrences of the three-node inter-DMN pathways that negatively correlated with visuospatial function. Occurrences of the 12 possible pathway' permutations are shown. A. The inter DMN-visual pathways. B. The inter DMN-Limbic pathways. C. The inter DMN-sensorimotor pathways.

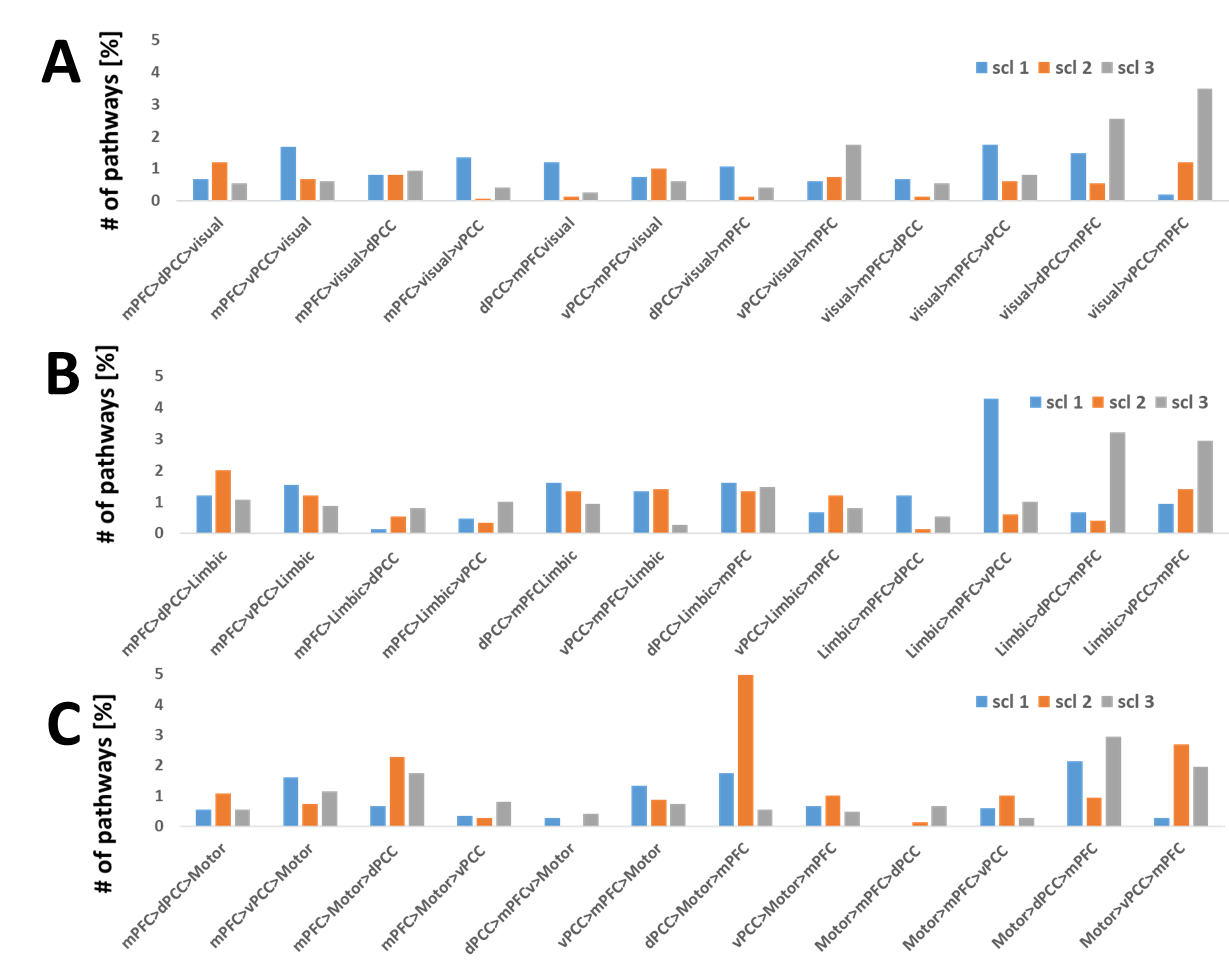

Figure 4: Occurrences of the three-node inter-DMN pathways that positively correlated with language. Occurrences of the 12 possible pathway' permutations are shown. A. The inter DMN-visual pathways. B. The inter DMN-Limbic pathways. C. The inter DMN-sensorimotor pathways.

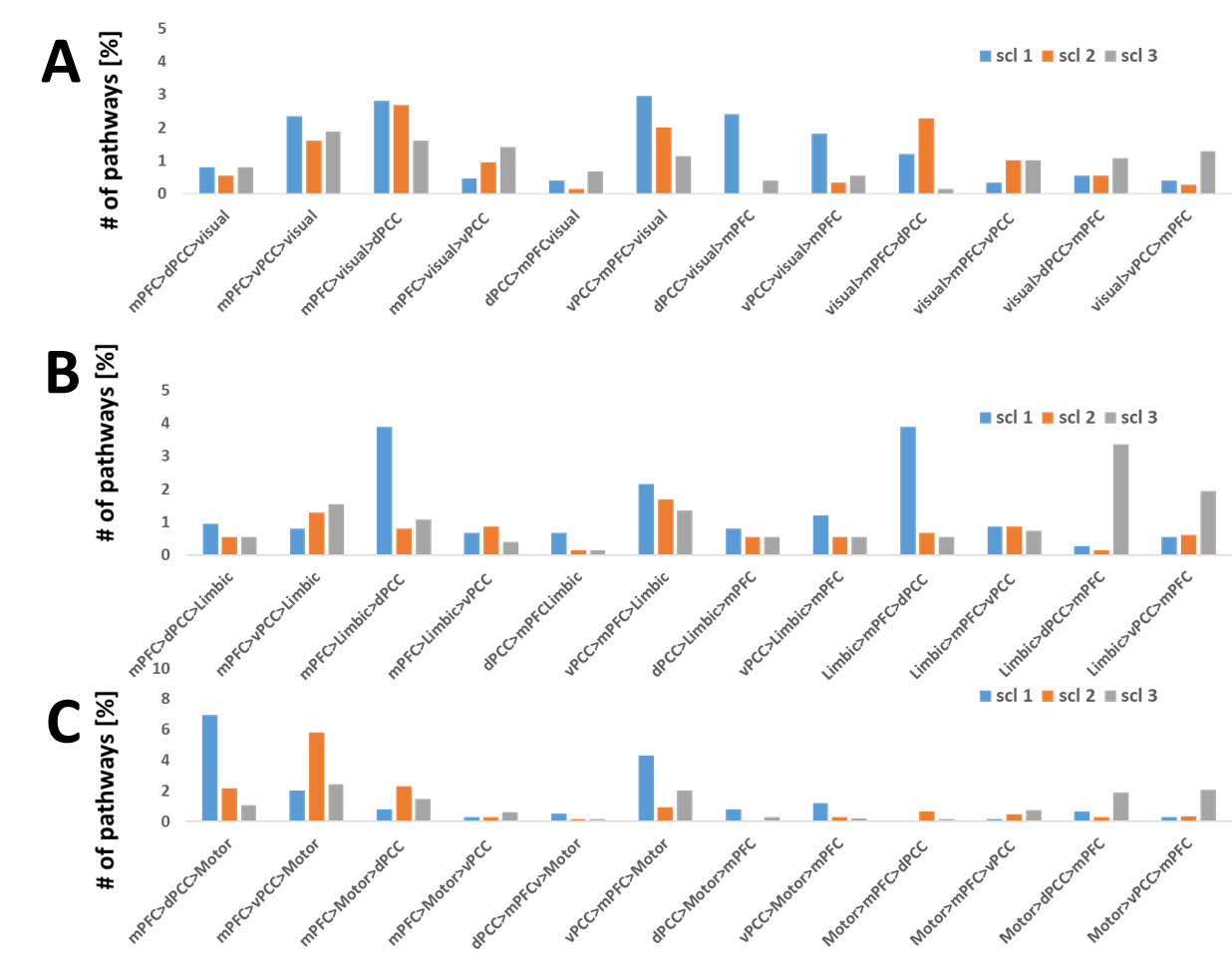

Figure 5: Occurrences of the three-node inter-DMN pathways that positively correlated with memory. Occurrences of the 12 possible pathway' permutations are shown. A. The inter DMN-visual pathways. B. The inter DMN-Limbic pathways. C. The inter DMN-sensorimotor pathways.
